# Varying thermal exposure, host-plant traits and oviposition behaviour across vegetation ecotones

**DOI:** 10.1101/2020.02.11.944439

**Authors:** Maria Vives-Ingla, Javier Sala-Garcia, Constantí Stefanescu, Josep Peñuelas, Jofre Carnicer

**Author notes:** corresponding authors Maria Vives-Ingla, Jofre Carnicer.

## Abstract

Vegetation cover generates local microclimatic gradients in the understorey, being especially pronounced at narrow ecotones linking open and forested habitats (*open–closed ecotones*). They provide key habitats for multiple insect communities and may largely determine the exposure of herbivorous insects to the increasing impacts of climate change. We report parallel measurements of microclimatic variables, multiple host-plant traits, and oviposition behaviour in Mediterranean populations of two *Pieris* butterflies across ecotones of vegetation cover. Open microhabitats were significantly warmer, drier, and more exposed to thermal amplification, which increased temperatures to values affecting insect larval survival. Host plants advanced their reproductive phenology and were shorter. Open microhabitats also inhibited the development of shade-adapted plants (e.g. *Alliaria petiolata*), decreasing fruit production. In contrast, the reproduction of sun-adapted host plants (e.g. *Lepidium draba*) was vigorous in the open microhabitats and completely inhibited in closed microhabitats, which were exclusively inhabited by non-reproductive ramets. Key plant traits for the selection of oviposition sites by butterflies, such as foliar water and chlorophyll contents, varied significantly across the open– closed ecotones. Foliar water content was always lower in the open microhabitats, whereas foliar chlorophyll gradients differed between sun- and shade-adapted plants. The oviposition behavior of *Pieris* butterflies across the ecotones differed significantly between the thermotolerant species (*P. rapae*, preferentially selecting open microhabitats) and the thermosensitive species (*P. napi*, selecting microhabitats protected by vegetation cover), matching the values of thermal susceptibility estimated from parallel heat tolerance assays of the populations. The larvae of the thermotolerant *Pieris* species grew under completely different thermal conditions due to differential microhabitat selection, indicating marked interspecific differences in thermal exposure (5–10 °C). These results suggest that the impacts of global warming in these communities will likely be mediated by open–closed ecotones, which determine pronounced local variability in thermal exposure, oviposition placement, and host-plant traits affecting larval performance in summer.

## 1. INTRODUCTION

The effects of climate change on natural systems have been consistently detected in many regions of the world and are predicted to increase as anthropogenic warming continues to intensify in the coming decades (Hoegh-Guldberg et al., 2018; IPCC, 2014; Parmesan, 2006; Parmesan & Yohe, 2003; Urban, 2015; Walther et al., 2002). Reported impacts of climate change on organisms, however, include a wide array of responses involving processes at multiple scales and levels of ecological organization (Moritz & Agudo, 2013; Parmesan, 2006; Scheffers et al., 2016). Climatic exposure in a habitat is due to the interaction of large-scale climatic conditions with site-specific geophysical attributes. Topography, vegetation structure, soil composition, and even surface roughness can locally modify macroclimatic conditions and generate a mosaic of microclimates (Bramer et al., 2018; De Frenne et al., 2013, 2019; Pincebourde, Murdock, Vickers, & Sears, 2016; Woods, Dillon, & Pincebourde, 2015). For example, local temperature gradients within a few meters can parallel gradients at larger, geographical scales (Lenoir et al., 2013; Pincebourde et al., 2016; Scherrer & Körner, 2010). Thermal variability in microhabitats is thus noteworthy, because the experience of climate by organisms ultimately depends on the way in which they sample these microclimatic mosaics (Bennett, Severns, Parmesan, & Singer, 2015; Pincebourde et al., 2016; Woods et al., 2015).

Microclimatic heterogeneity contributed to the occurrence of microrefugia in the past (Dobrowski, 2011) and may similarly play a key role in mediating the effects of current climate change on ecological systems, as several studies have already suggested (Bennett et al., 2015; Bonebrake, Boggs, Stamberger, Deutsch, & Ehrlich, 2014; Carnicer et al., 2019, 2017; De Frenne et al., 2013; Hindle, Kerr, Richards, & Willis, 2015; Kearney, Shine, & Porter, 2009; Lenoir et al., 2013; Pincebourde et al., 2016; Scherrer & Körner, 2010; Suggitt et al., 2018; Sunday et al., 2014; Woods et al., 2015). For example, De Frenne et al. (2013) recently found that an increase in warm-adapted plant species in the understorey of temperate forests of the northern hemisphere was being attenuated in forests whose canopies had become denser. This result was attributed to the buffering against the impacts of macroclimatic warming provided by canopy closure, lowering ground-layer temperatures and increasing relative air humidity and shade. In addition to acting as a buffer for the understorey, forest cover can also protect insect communities that rely on these host plants. Limited thermal buffering of vegetation in the Mediterranean biome was identified as a key factor exacerbating the decline of a population of a drought-sensitive species (Carnicer et al., 2019). Other interacting negative factors included the reduction of host-plant quality due to the seasonal progression of plant phenology, the amplification of foliar temperatures linked to reduced plant transpiration, and increasing impacts of summer drought at the multidecadal scale (Carnicer et al., 2019). Detailed descriptions of the effects of vegetation-cover gradients on multiple host-plant traits and thermal microconditions, however, are currently lacking for most interactions between plants and animals.

We determined whether local spatial gradients in vegetation cover (hereafter open–closed ecotones) induced microclimatic heterogeneity and plasticity of host-plant traits related to plant quality for insect hosting and herbivory. We also assessed whether microhabitat variation could be associated with different oviposition behaviour of butterflies. The selection of oviposition sites may have important implications for offspring survival and performance by strongly influencing the environment and resource availability before and after hatching (Gibbs & Van Dyck, 2009). Numerous factors can alter oviposition behaviour, such as temperature, the state and distribution of host plants, and the surrounding vegetation (Gibbs & Van Dyck, 2009). For example, temperature can influence the selection of microhabitats to lay eggs by directly enhancing oviposition and/or by modifying the amount of time and the number of suitable locations that are available for egg-laying. The development of eggs and larvae can be also affected by microhabitat conditions. In addition to directly affecting growth, microclimatic variation can also induce plastic shifts in host-plant quality (Merckx, Serruys, & Van Dyck, 2015), which can synergistically act on larval development (Bauerfeind & Fischer, 2013). All these processes have been extensively studied but have usually been treated separately or only a few habitat variables and/or host-plant traits have been considered (see Gibbs & Van Dyck, 2009 for a review). Integrative studies with comprehensive, parallel measurements of multiple host-plant traits, oviposition behaviour, and microclimatic variables across local spatial gradients are thus warranted. We studied the influence of open–closed ecotones on two host plants, *Alliaria petiolata* (M. Bieb.) Cavara & Grande and *Lepidium draba* L., in two Mediterranean sites, both harbouring populations of the butterflies *Pieris napi* L. 1758 and *P. rapae* L. 1758 to integrate these processes in a single study and to describe their simultaneous variation. More specifically, we quantified the seasonal dynamics (objective *i*) and the spatial variation across open–closed ecotones (objective *ii*) of key microhabitat variables:

a. microclimatic conditions where the two plant species that host *P. napi* and *P. rapae* grow,
b. host-plant phenology and reproductive output, and
c. host-plant morphological and physiological traits influencing butterfly oviposition.

We then determined whether the patterns of spatial variation were maintained across seasons and host-plant phenological stages (objective *iii*). Finally, in objective *iv* we tested whether the two *Pieris* species, which differ in habitat affiliations, selected different microhabitats from open–closed ecotones to oviposit and had different larval thermal susceptibilities.

## 2. MATERIALS & METHODS

### a. Study system

We studied two cohorts of the host plants *A. petiolata* and *L. draba* to quantify the effects of vegetation cover on the variation of local microclimatic conditions and host-plant traits. The two species were distributed at two different sites (*A. petiolata* at site 1 and *L. draba* at site 2) belonging to different protected areas of the north-eastern Iberian Peninsula, 50 km from each other (Figs. S1 and S2). Both sites are along two transects that have long been monitored by the Catalan Butterfly Monitoring Scheme (CBMS, Pollard & Yates, 1993; Stefanescu, 2000) and contain abundant and intensively studied populations of the butterflies *P. napi* and *P. rapae* (Carnicer et al., 2019). Site 1 is in a mid-elevation area (539 m a.s.l.; Can Jordà, La Garrotxa Volcanic Zone Natural Park) populated by Mediterranean, sub-Mediterranean, and Eurosiberian vegetation. Site 1 is a heterogeneous landscape of evergreen and deciduous woodlands (including holm oak, deciduous oaks, and beech as the main arboreal species), meadows, pastures, arable land, and natural ponds. Site 2 is in a lowland coastal wetland (Aiguamolls de l’Empordà Natural Park) surrounded by riparian deciduous forests, bush and bramble thickets, reed beds, and irrigated cropland.

We studied the oviposition behaviour of closely related *P. napi* and *P. rapae.* The green-veined white butterfly (*P. napi*) is a Holarctic species tightly linked to humid habitats. It can be found throughout Catalonia, except in the driest areas (Vila, Stefanescu, & Sesma, 2018). It is more locally distributed in lowlands, and declining trends among these populations have been associated with the increasing impacts of summer drought (Carnicer et al., 2019). The small white butterfly (*P. rapae*) is a more generalist, thermophilous, and ubiquitous species. It is very common in agricultural and ruderal areas, and its populations tend to be stable or slightly increasing (Vila et al., 2018). Both species lay individual eggs on Brassicaceae species. *Pieris rapae* uses a greater diversity of host plants (both natural and cultivated), but *P. napi* restricts its oviposition to several crucifers common in humid habitats (e.g. *A. petiolata, Brassica nigra* (L.) W. D. J. Koch, *L. draba*, and *Cardamine* spp.; Vila et al., 2018; Carnicer et al. 2019).

*A. petiolata* and *L. draba* are both used as host plants by the two butterfly species and are dominant at their study sites. *Alliaria petiolata* is a biennial herb adapted to shade and is thus common in damp soils at the edges of deciduous forests (de Bolós & Vigo, 1990) but can also grow in highly contrasted environmental conditions, exhibiting considerable plasticity in different habitats (Cavers, Heagy, & Kokron, 1979). It has heart-shaped leaves 2–20 cm in length. Seedlings emerge during spring (even from late spring to early summer) and persist as rosettes throughout the first year, until the next growing season when inflorescences are initiated. Fruits are ascendant siliques 20–70 mm long. We studied *L. draba*, which is a perennial, rhizomatous, and sun-adapted herb usually found in ruderal areas and field margins with deep soil, in the lowlands (site 2) (de Bolós & Vigo, 1990). Its leaves are ovate, about 1.5– 10 cm long. Flowers are grouped in corymbs and produce indehiscent silicules. Its extensive, multi-branched rhizomes are notably capable of producing many new shoots, which can develop into large monocultural stands (Francis & Warwick, 2008).

### b. Microenvironmental and host-plant variation

The landscape mosaic of the two study sites was characterised by spatial gradients of vegetation cover between open and closed microhabitats and their transition zones (*open–closed ecotones*). We monitored cohorts of 152 individuals of *A. petiolata* and 353 individuals of *L. draba* distributed across the ecotones. Each individual was assigned to one of four categories of microhabitats for assessing the influence of vegetation cover on the variabilities of the microclimates and host-plant traits: open (O), semi-open (SO), semi-closed (SC), and closed (C). These categories were based on detailed measurements of the dynamics of vegetal cover conducted at the sites (Text S1 and Figs. S2 and S3). Both cohorts included individuals from the four types of microhabitats.

We continuously monitored 19 host-plant traits and microclimatic conditions in the *A. petiolata* and *L. draba* cohorts to quantify their seasonal variation (Table 1). We randomly selected 12 host plants for each cohort each monitoring day, with three samples for each category of microhabitat (O, SO, SC, and C). The selection procedure ensured that plants were randomly chosen without repetition to avoid pseudoreplication. A representative basal, medial, and apical leaf was chosen for each plant, and its state (green/senescent) was recorded. Microclimatic and host-plant measurements were conducted from March to October 2017, repeating the same sampling protocol every 15 days in each microhabitat.

**Table 1.**
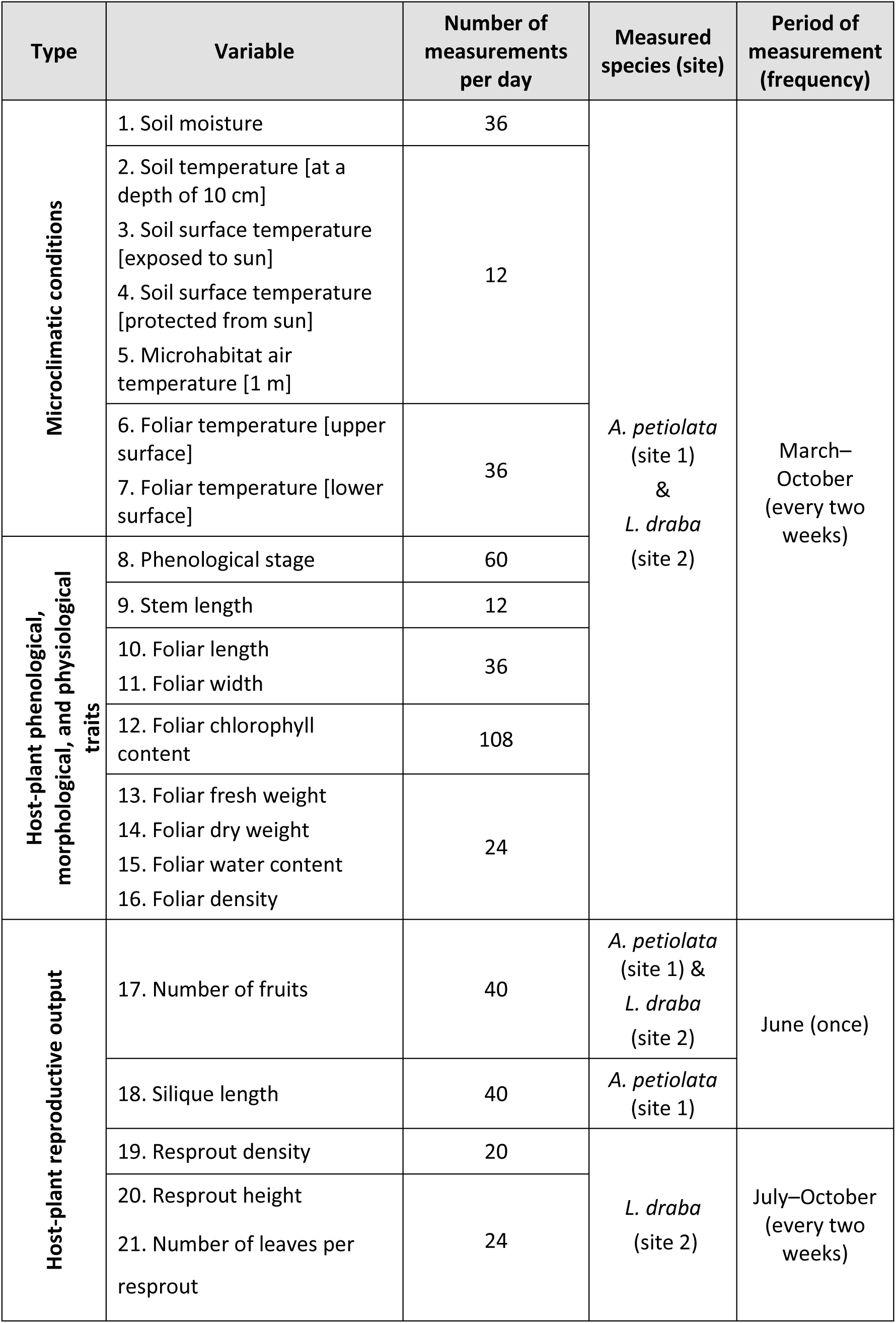
Summary of the 21 variables measured during microclimatic and host-plant monitoring.

The volumetric water content of the soil (% by volume, Table 1 variable 1) was measured at three points near each plant using a DELTA-T SM150 (Delta-T Devices Ltd, Cambridge, UK) soil-moisture sensor kit. A penetration thermometer (HANNA HI98509, Hanna Instruments Ltd, Eibar, Spain) was used for measuring soil temperature at a depth of 10 cm (Table 1 variable 2). Soil surface temperature (Table 1 variables 3, 4), microhabitat air temperature (Table 1 variable 5), and foliar surface temperature (Table 1 variables 6, 7) were measured using a wire K-type thermocouple probe (Omega SC-TT-KI-30-1M, Omega Engineering Ltd, Manchester, UK) attached to a hand-held thermocouple thermometer (Omega HH503, Omega Engineering Ltd, Manchester, UK, and HANNA HI935005N, Hanna Instruments Ltd, Eibar, Spain). Average measurements (at least three records) were kept. The temperatures were measured between 10:00 and 16:00, and the time and wind and radiation conditions were recorded. Soil surface temperature was measured near the host plants, replicating it in areas exposed to direct solar radiation and in shaded areas. Air temperature was measured at a height of 1 m immediately above the host plant. Foliar temperature was measured on the upper and lower surfaces. We calculated foliar thermal amplification as the difference between foliar temperature and the maximum recorded environmental temperature of the corresponding day to compare foliar thermal microconditions with standard measurements of the local weather. Daily records of environmental temperature were obtained from two meteorological stations near the study sites and within the same elevational range (Fig. S1). Additionally, microclimatic conditions were continuously recorded with standalone data loggers (Lascar Electronics EL-USB-2-LCD, Salisbury, UK). Eight data loggers were placed 25 cm above the soil near the host plants in each microhabitat type and site (see below). The sensors were programmed to measure temperature (°C) and relative humidity (RH, %) hourly.

We assessed plant phenological status (Table 1 variable 8) by classifying the individuals in one of four phenological stages: early vegetative (spring rosettes and young shoots before budding), reproductive (plants with buds, flowers, and/or fruits), senescent, and late vegetative (late *A. petiolata* seedlings and *L. draba* resprouts emerging in summer). The length, width, and chlorophyll content (Table 1 variables 10–12) of each leaf were measured. Chlorophyll content was estimated as the mean of three measurements from a MINOLTA SPAD-502 (Konica Minolta Sensing, Valencia, Spain) chlorophyll meter. Finally, leaves were severed and immediately weighed (fresh weight, FW; Table 1 variable 13) using a Pesola PJS020 Digital Scale (PESOLA Präzisionswaagen AG, Schindellegi, Switzerland) for calculating water content. The leaves were oven-dried in the laboratory at 60 °C for two days to a stable weight (dry weight, DW; Table 1 variable 14). The ratio of foliar water content (to DW, Table 1 variable 15) was defined as (FW-DW)/DW. The ratio DW/foliar length was calculated as a proxy for foliar density (Table 1 variable 16).

### c. Host-plant reproductive output

Fruit production was measured in each microhabitat type as an indicator of differential host-plant fitness. A minimum of seven individuals were sampled for each microhabitat type. The number of fruits (siliques for *A. petiolata* and silicules for *L. draba*) per plant was counted (Table 1 variable 17). For *A. petiolata*, we also measured host-plant height and silique length (Table 1 variable 18). For *L. draba*, we additionally conducted a census of newly emerging resprouts. The density of resprouts was quantified beginning in July when the first shoots emerged from resprouting rhizomes (Table 1 variable 19). Five 25-cm quadrats were randomly placed in each microhabitat type. The total number of resprouts per unit area were counted, and three resprouts were then randomly selected for measuring their heights (Table 1 variable 20) and counting their total numbers of leaves (Table 1 variable 21).

### d. Oviposition behaviour

As previously stated, *P. napi* and *P. rapae* are usually associated with different habitat types (*P. napi* with humid areas and *P. rapae* with open and dry areas). Microhabitat use by these species and the interactions with vegetation structure nevertheless remain poorly described and may vary depending on the kind of behaviour (e.g. basking sites do not coincide with oviposition sites) and on the time of day and season (Dennis, 2004). We assessed whether the differences in broad habitat preferences between the species led to different microhabitat selections for oviposition across open–closed ecotones. We tested this hypothesis by carrying out censuses of oviposition behaviour at the two study sites. Females were followed for replicated periods of 45 min to count the number of eggs they laid and record the microenvironmental conditions. The censuses fully covered the entire daily period of flight activity, between 9:00 and 19:00, and were conducted in summer 2017 (lowland site, two days) and 2018 (mid-elevation site, four days). They were simultaneously performed in the various microhabitat types, carefully balancing the time spent in each type. Oviposition was considered to occur when females that landed on a leaf were observed to curl their abdomen and remain in this position for at least three seconds. Species, hour, egg position (upper vs lower surface of leaves), and microhabitat type were recorded. The temperature of the leaves where eggs were laid was also recorded when possible using a thermocouple (see previous sections) immediately after the female left the plant. Additional ovipositions during host-plant monitoring were also integrated into the final data set. Table S1 contains a summary of the census variables and the number of females and eggs. Mean temperature between 13:00 and 17:00 (when 85% of ovipositions occurred) recorded with the data loggers from June to September was calculated for each microhabitat type and site to assess the thermal conditions during oviposition. We also assessed mean thermal amplification, defined as the difference between the mean daily temperatures from the microclimatic data loggers and the temperatures from standardised local meteorological stations.

### e. Ecophysiological assays of heat tolerance

If as hypothesized *P. napi* oviposits in more closed and buffered microhabitats than does *P. rapae, P. napi* larvae may be more susceptible than *P. rapae* larvae to thermal stress. Heat tolerance assays determining the time to thermal death can predict from first principles how thermal stress (depending on both its intensity and duration) can affect larval survival (Deutsch et al., 2008; Rezende, Castañeda, & Santos, 2014). We implemented a static heat tolerance experiment using *P. napi* and *P. rapae* larvae to determine whether the two species had different survival responses to thermal stress. Text S2 provides a more detailed description of the experiments.

### f. Data analyses

All data were analysed using R 3.6.1 (R Core Team, 2019). We used local polynomial regressions between each variable and Julian day to assess the temporal dynamics of the microenvironmental conditions and host-plant traits (objective *i*) (Table S2). The regression fit was applied separately to each microhabitat type (O, SO, SC, and C) at each site. The trends for the host-plant variables were grouped by plant developmental stage (i.e. flowering spring plants vs newly emerged or non-flowering summer individuals). Weekly abundances of *P. napi* and *P. rapae* were obtained from the CBMS transects at both sites (2017). An index of butterfly abundance for each recording day was calculated as the number of counts divided by the length of the transect (in km). A local polynomial regression analysis against Julian day (neighbourhood parameter α = 0.25) was then applied to determine the phenologies of both butterflies at both sites.

Each variable was modelled against microhabitat type for describing the variation of microenvironmental conditions and host-plant traits across open–closed ecotones at each site (objectives *ii* and *iii*). An ANOVA was applied followed by a post-hoc Tukey HSD test calculated using the *emmeans* package (Lenth, 2019). The analyses were performed for the entire sampling period (objective *ii*) and for specific phenological stages and seasons (objective *iii*). We quantified the date for the onset of flowering to assess phenological differences across the open–closed ecotones. Changes in the daily proportion of individuals in each microhabitat type that were at their reproductive stage were assessed using the data from the phenological censuses (Table 1 variable 8). A prediction curve and its 95% confidence interval were obtained from the fit of a local polynomial regression between the proportion of reproductive individuals and Julian day (neighbourhood parameter α = 0.5, Table S3). The onset of flowering for each microhabitat type and site was calculated as the first Julian day when 50% of the plants were flowering.

We applied generalised linear mixed models (GLMMs) to determine whether the two butterfly species selected different microhabitats to oviposit (objective *iv*) (Bolker et al., 2009; Zuur, Ieno, Walker, Saveliev, & Smith, 2009). The number of females ovipositing was used as the response variable. The type of microhabitat, the species, their interaction, and census duration were added as fixed factors, and Julian day and time of the census (9:00–11:00, 11:00–13:00, 13:00– 15:00, 15:00–17:00, and 17:00–19:00) were treated as random factors. The model was fitted using the bglmer function of the *blme* package (Chung, Rabe-Hesketh, Dorie, Gelman, & Liu, 2013) by maximum likelihood (Laplace approximation), a Poisson error distribution with a log link function and imposing zero-mean normal priors on the fixed effects to avoid complete separation (Bolker, 2015). p values for the fixed effects were calculated by parametric bootstrapping using the *afex* package (Singmann, Bolker, Westfall, & Aust, 2019). Finally, the predicted distribution of ovipositing females during a day was also estimated from the conditional modes of time grouping factor. Data from both sites were compiled into a single data set and treated together, because they had similar distributions of eggs across the open– closed ecotones. The number of eggs was not used as a response variable to avoid pseudoreplication linked to differences in oviposition behaviour between the species (e.g. *P. napi* lays more eggs than *P. rapae* before switching to new hosts, Friberg & Wiklund, 2019).

## 3. RESULTS

### a. Seasonal dynamics of microclimatic conditions and host-plant traits (objective *i*)

The annual cycles of variation in microclimatic conditions, host-plant traits, and butterfly phenologies are shown in Fig. 1, highlighting the main differences between the C and O microhabitats. The thermal variables (Table 1 variables 2–7) were strongly correlated (*p* = 0.0042 ± 0.0198 and *R*^2^ = 0.97 ± 0.04 in the pairwise correlations between all thermal variables for each site and microhabitat type). Temperatures were higher in the O than C microhabitats at both sites. The difference was especially pronounced at the soil surface, where soils in the O microhabitats were a mean of 10.1 ± 6.9 °C warmer (Fig. 1A, B) and drier (Fig. 1C, D). Soil humidity in the O microhabitats markedly decreased as summer temperatures increased, but the trends of soil humidity were more stable in the C microhabitats.

**Figure 1.**
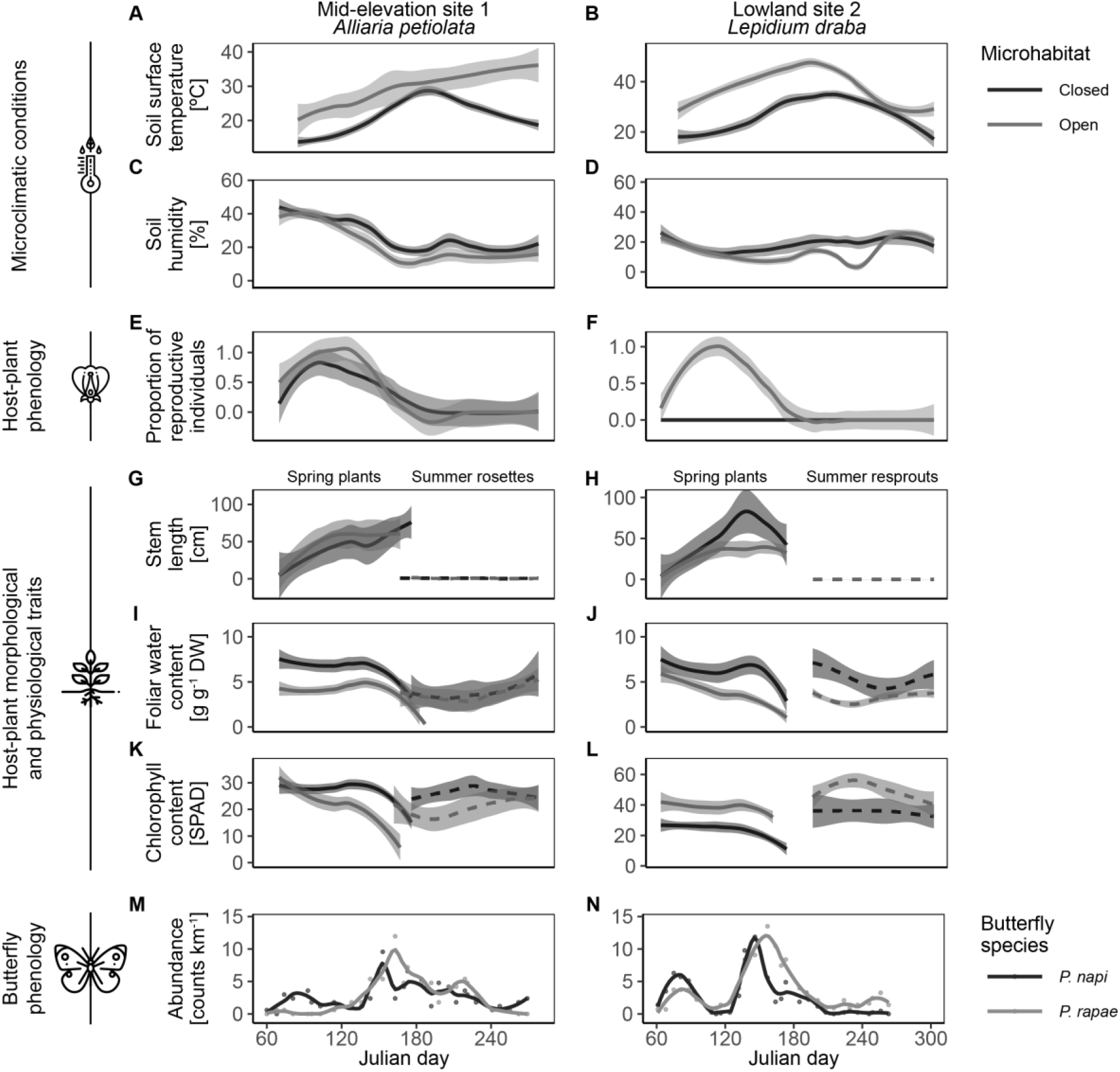
Seasonal patterns of variation of microclimatic conditions (A–D), host-plant phenological (E and F) and morphological and physiological traits (G–L), and butterfly phenology (M and N). The assessment of host-plant morphological and physiological traits (G–L) separated individuals into those that emerged and developed in early spring (spring plants) and those that emerged since June (summer rosettes and resprouts). Shaded areas indicate the 95% confidence interval around each trend, except for the soil surface temperature, where the shaded areas represent the 50% confidence intervals. A summary of the fits of local polynomial regressions between the response variables and Julian day is presented in Table S2.

Individuals of both host-plant species began to reproduce earlier in spring and in larger proportions in the O than the C microhabitats (Figs. 1E, F and S4). More closed (C and SC) microhabitats completely inhibited the onset of flowering of the sun-adapted species, *L. draba*, at the lowland site (Figs. 1F and S4). Consistent with these phenological observations, total stem length was stabilized earlier in the O microhabitats, leading to shorter mature plants (Fig. 1G, H). Foliar water and chlorophyll contents decreased in both host plants (Fig. 1I–L) as they senesced after fructification during late spring and early summer (Julian days 140–180) (Figs. 1E, F and S4). Only non-flowering first-year rosettes (*A. petiolata*) and summer rhizome resprouts (*L. draba*) remained in midsummer after senescence (Fig. 1G, H). First-year rosettes of *A. petiolata* notably coexisted in June with the reproductive stage of second-year individuals, which remained green (Figs. 1G, I, K and S4). In contrast, there was no temporal overlap between reproductive *L. draba* plants and new summer resprouts, which did not appear until mid-July. The complete senescence of *L. draba* individuals in late June thus led to a period of scarcity of fresh host plants for 2–3 weeks until the emergence of midsummer resprouts in mid-July (Figs. 1H, J, L and S4). The abundances of the two *Pieris* butterfly species peaked at both sites when the dominant host plants were experiencing these changes in late spring and early summer (Fig. 1M, N). The abundance of the next butterfly generations at the lowland site decreased sharply after the peak, coinciding with the period of *L. draba* scarcity (Fig. 1N).

### b. Microclimatic and host-plant variation across open–closed ecotones (objective *ii*)

The variation of the microclimates and host-plant traits were significantly associated with the open–closed ecotones (Fig. 2 and Table S4). Temperatures were significantly higher in the O and SO than the C and SC microhabitats, as suggested in the previous section. Upper foliar temperatures averaged 5 °C higher in the O microhabitats, reaching or exceeding the maximum environmental thermal records (Fig. 2A–D). Foliar thermal amplification was more pronounced at the lowland site than in mid-elevation areas (Fig. 2C, D), even though environmental temperatures were lower at the lowland than the mid-elevation site (Fig. 2A, B; note that the grey shaded areas in the panels depict the distribution of maximum daily environmental temperatures). Foliar temperature at the lowland site reached 40 °C, surpassing values that could affect the survival of *Pieris* larvae, based on the analyses of the heat tolerance experiments (TE6h in Figs. 2B and S5 and Text S2). In sharp contrast, differences between foliar temperatures and maximum daily records were ≤0 °C at the mid-elevation site (Fig. 2C), where critical thermal limits were rarely surpassed (TE6h in Fig. 2A). Soils were significantly drier in the O microhabitats (Fig. 2E, F). The plants flowered earlier in the O microhabitats (Fig. 2G, H), consistent with the reported differences in microclimatic conditions and phenological trends analysed in Figs. 1E, F and S4. The host plants in these microhabitats were significantly shorter (Fig. 2I, J), with smaller leaves (Fig. 2K, L) that had lower ratios of water content (i.e. less water mg^-1^ foliar DW, Fig. 2M, N). This was accompanied by denser leaves at the lowland but not at the mid-elevation site, where foliar density was highest in the SO microhabitats (Fig. 2O, P). Foliar chlorophyll content had opposite spatial patterns at the two sites (Fig. 2Q, R). Chlorophyll content for *L. draba*, the dominant and sun-adapted host plant at the lowland site, was highest in the O microhabitat and lowest in the C microhabitat (Fig. 2R). On the contrary, chlorophyll content for *A. petiolata* (shade-adapted) at the mid-elevation site was lowest in the most open microhabitat (Fig. 2Q).

**Figure 2.**
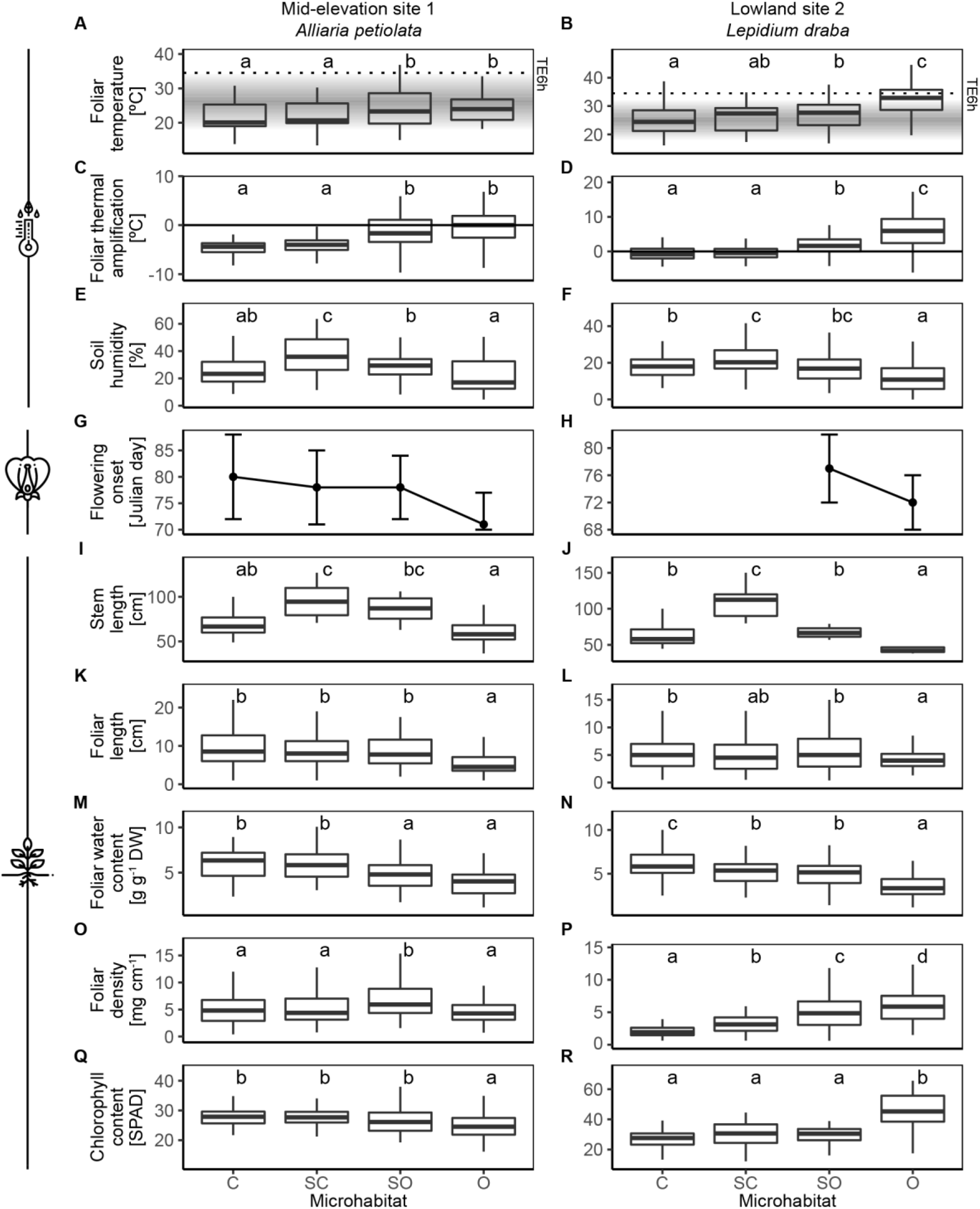
Spatial patterns of variation of microclimatic conditions (A–F) and host-plant phenological (G and H) and morphological and physiological traits (I–R) across the open–closed ecotones. Different letters indicate significant differences of the response variable between the microhabitat types in the Tukey HSD test. The grey shaded areas in panels A and B correspond to the maximum daily temperatures recorded by the local meteorological stations. The dotted lines correspond to the thermal thresholds that would lead to larval death with a daily exposure of six hours during the entire developmental period (TE6H) estimated by the heat tolerance experiments. The horizontal lines in panels C and D indicate foliar temperatures equal to maximum daily temperature. Positive values correspond to foliar thermal amplification, and negative values imply thermal buffering. A summary of the results is presented in Table S4. C, closed microhabitat; SC, semi-closed micrhabitat; SO, semi-open microhabitat; and O, open microhabitat.

Variation of fruit production across the open–closed ecotones closely paralleled the contrasting patterns of foliar chlorophyll content between host plants (Fig. 3A, B). Fruit production for *A. petiolata* was highest in the SO, SC, and C microhabitats and was much lower in the O microhabitats (Fig. 3A). In contrast, sexual reproduction for *L. draba* was completely inhibited in the C and SC microhabitats, but fruit production was high in the O and SO microhabitats (Fig. 3B). The vegetative production of summer resprouts of *L. draba* was significantly affected by the conditions in the open–closed ecotones, showing the highest density of resprouts and total number of leaves in the O microhabitats (Fig. 3C, D).

**Figure 3.**
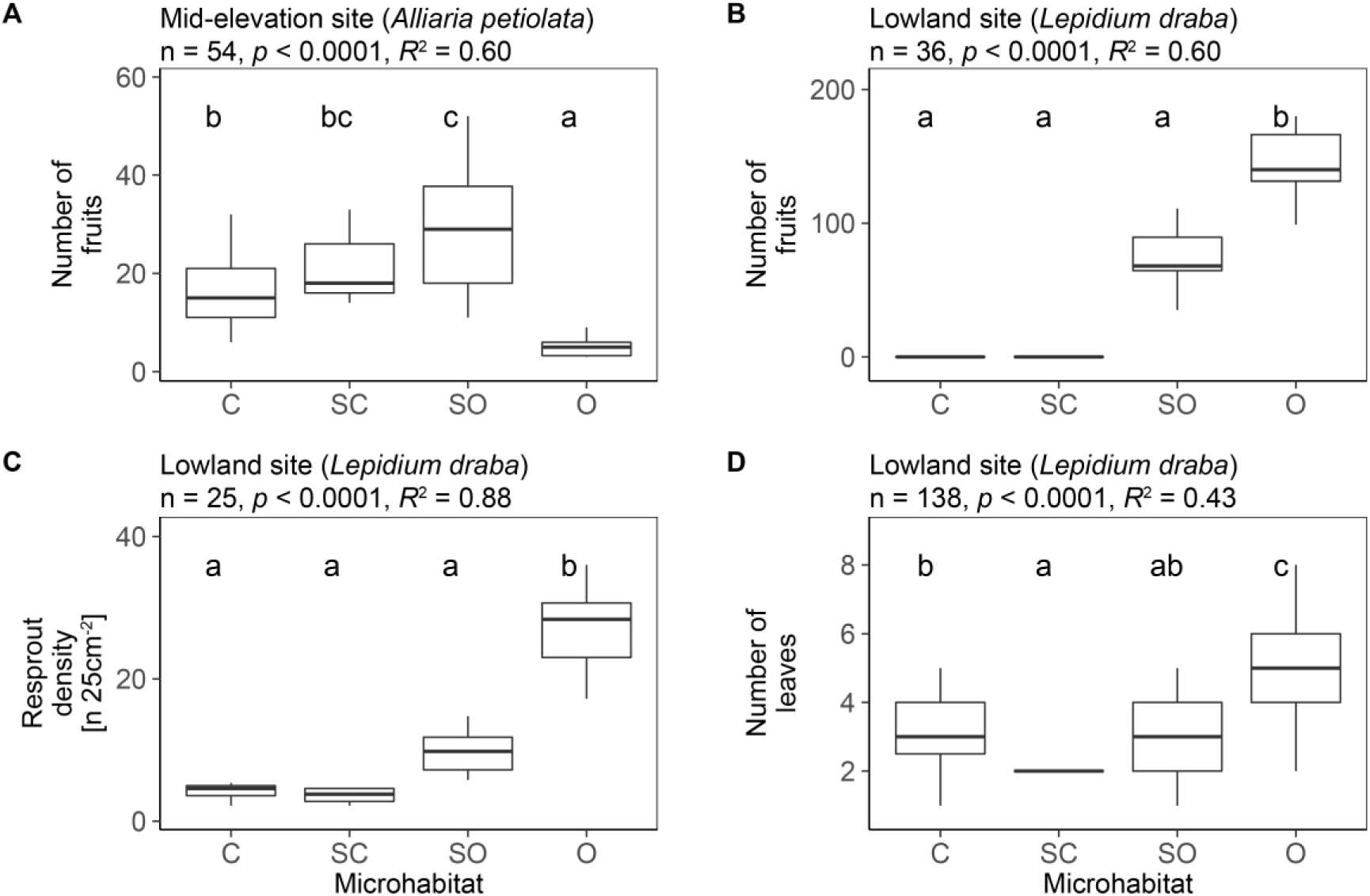
Variation in the reproductive output of the host plants across the open–closed ecotones. Different letters indicate significant differences of the response variable between the microhabitat types in the Tukey HSD test. C, closed microhabitat; SC, semi-closed microhabitat; SO, semi-open microhabitat; and O, open microhabitat.

### c. Seasonal variation of the spatial patterns across the open–closed ecotones (objective *iii*)

The microclimatic differences between the ecotone microhabitats remained significant across seasons (from spring to autumn) (Figs. 4, S6, S7 and Table S5). Temperatures at both sites were significantly higher in the O and SO microhabitats in all seasons (Fig. 4A–H). Foliar temperatures for *A. petiolata* in the C and SC microhabitats were particularly buffered relative to the highest environmental temperatures in summer (Fig. 4I–L). In contrast, foliar temperatures for *L. draba* were significantly amplified relative to maximum environmental temperatures, especially in the O and SO microhabitats (Fig. 4M–P). Foliar thermal amplification led temperatures to sufficiently high values to affect larval survival in all seasons in the O microhabitats at the lowland site (Fig. 4E–H), whereas this critical thermal limit was only surpassed in midsummer at the mid-elevation site (Fig. 4C). Overall, the open–closed ecotone structure largely determined the thermal buffering and amplification processes that operated in the host-plant microhabitats, modifying their thermal exposure.

**Figure 4.**
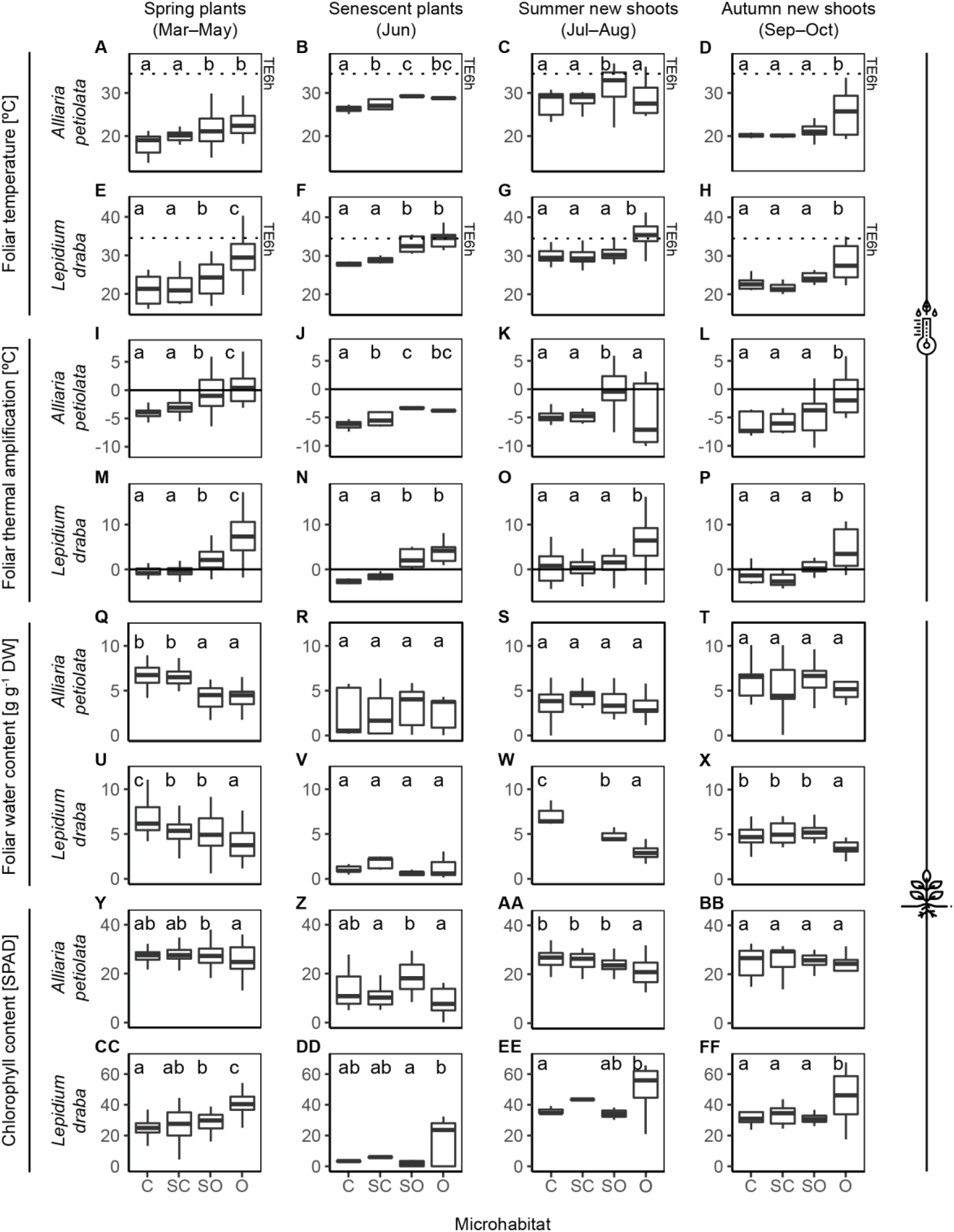
Seasonal variation of the spatial patterns of variation across open–closed ecotones. A–P, microclimatic conditions; Q–FF, host-plant traits. Different letters indicate significant differences of the response variable between the microhabitat types in the Tukey HSD test. The dotted lines in panels A–H correspond to the thermal thresholds that would lead to larval death with a daily exposure of six hours during the entire developmental period (TE6H) estimated by the heat tolerance experiments. The horizontal solid lines in panels I–P indicate foliar temperatures equal to the maximum daily temperature. Positive values correspond to foliar thermal amplification, and negative values imply thermal buffering. No foliar sample was weighted (panel W), and foliar chlorophyll content was not measured in one case (panel EE), due to the small number and size of midsummer *Lepidium draba* resprouts in the semi-closed microhabitat. A summary of the results is presented in Table S5. C, closed microhabitat; SC, semi-closed microhabitat; SO, semi-open microhabitat; and O, open microhabitat.

Foliar water and chlorophyll contents, which are key plant traits for the selection of butterfly oviposition sites (Gibbs & Van Dyck, 2009; Myers, 1985; Stefanescu, Peñuelas, Sardans, & Filella, 2006; Wolfson, 1980), also varied significantly across the open–closed ecotones (Fig. 4Q–FF). Their spatial variation was especially pronounced in spring plants of both host-plant species (Fig. 4Q, U, Y, CC). As previously stated, foliar water content was similar in both host-plant species (i.e. lower spring contents in the O and SO microhabitats, Fig. 4Q, U), whereas foliar chlorophyll content varied in opposite directions (i.e. contents in O microhabitats were lowest for *A. petiolata* but highest for *L. draba*, Fig. 4Y, CC). The decreases in foliar water and chlorophyll contents during plant senescence were larger for *L. draba*.

### d. Microhabitat selection during oviposition for the two *Pieris* species in the open– closed ecotones (objective *iv*)

Both *P. napi* and *P. rapae* distributed their eggs unequally across the open–closed ecotones. Furthermore, the microhabitats selected by females for oviposition differed significantly between the two butterfly species (microhabitat × species p = 0.0002, GLMM analysis; Table 2). *Pieris napi* laid more eggs on host plants distributed across the SO and SC microhabitats (jointly termed *intermediate open–closed microhabitats*, hereafter OC). In sharp contrast, *P. rapae* mainly laid eggs in the O microhabitats (Fig. 5A, B). Temperatures during oviposition were significantly lower in the OC microhabitats than in open areas (Fig. 5C, D), where thermal amplification was more severe (Fig. 5E, F). Foliar temperature during oviposition was accordingly higher in the leaves selected by *P. rapae* (Fig. 5G). Ovipositing females were predicted to be more numerous between 13:00 and 15:00 (Fig. 5H). Consistent with these results, the heat tolerance experiments indicated that *P. rapae* was more thermotolerant (i.e. their larvae were more tolerant of intense thermal stresses, Fig. S5 and Text S2).

**Table 2.** Generalized linear mixed model of the number of ovipositing female butterflies. Results from the parametric bootstrap test of the fixed effects.

| Fixed effect | $\chi^2$ | df | p |
| --- | --- | --- | --- |
| Microhabitat type | 9.27 | 2 | 0.0097 |
| Species | 0.19 | 1 | 0.6652 |
| Census duration | 0.67 | 1 | 0.4131 |
| Microhabitat type × species | 17.14 | 2 | 0.0002 |

**Figure 5.**
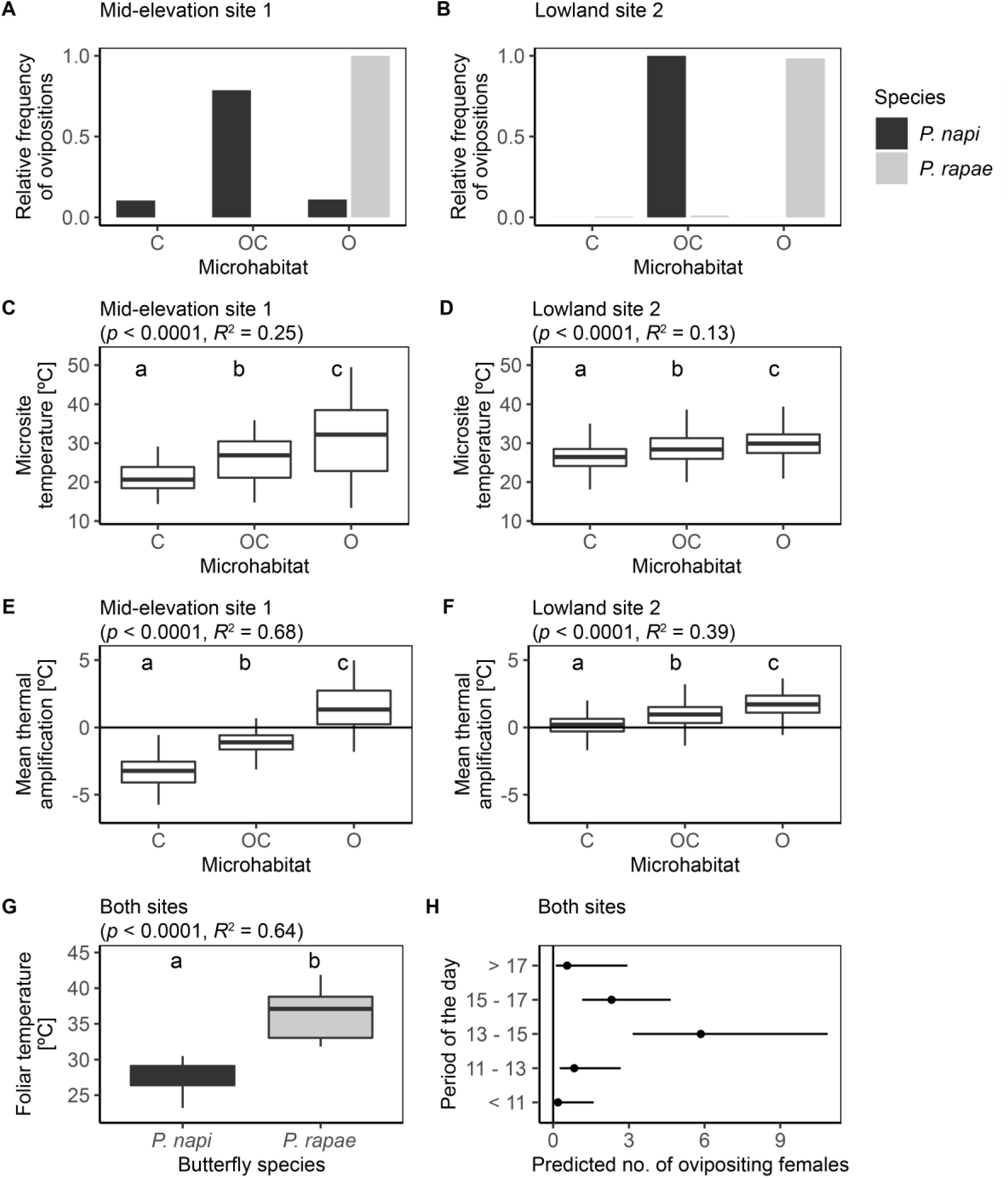
Results of the census of oviposition. A and B: distribution of the relative number of ovipositions for each species across the open–closed ecotones. C and D: mean summer temperature during oviposition (13:00–17:00) recorded by the standalone data loggers. E and F: difference between daily mean temperatures from the records of the microenvironmental data loggers and the local meteorological stations. G: foliar temperature of the lower surface during oviposition. H: predicted distribution of ovipositing females during the day in the GLMM.

## 4. DISCUSSION

Forest development is a key element of landscape engineering by the modification of surrounding physical conditions and resource abundance (Jones, Lawton, & Shachak, 1994). Our results indicated that multiple biotic and abiotic parameters, such as microclimatic heterogeneity, understorey host-plant variation, and butterfly ovipositing behaviour, were structured paralleling the gradients of vegetation cover across the open–closed ecotones. The O and SO microhabitats were significantly warmer, drier, and more exposed to thermal amplification that could elevate temperatures to values affecting larval survival (Figs. 1A–D, 2A– F, 4A–P, and S5). In contrast, C microhabitats benefitted from the buffering provided by the canopy cover, especially at the mid-elevation site 1 where the forest was more developed. Temperatures were thus lower, soil humidity was higher, and temporal dynamics were smoother there. Variation in the microhabitat structure across ecotones not only affected microclimatic conditions but also influenced all host-plant traits. The reproductive phenology of the host plants was advanced in the O microhabitats, and the plants were shorter and had smaller leaves with lower water contents (Figs. 1E–J, 2G–N, and 4Q–X). Other traits such as foliar chlorophyll content and reproductive output, however, had opposite patterns of spatial variation between host-plant species, indicating different shade-adaptation strategies. O microhabitats clearly inhibited the development of *A. petiolata* (shade-adapted), where the phenotype was less vigorous, with lower chlorophyll content and fruit production. In contrast, *L. draba* (sun-adapted) could not mature sexually and produced fewer resprouts in the C microhabitats (Fig. 3).

Microhabitat structure also differentially influenced the selection of oviposition sites by the two sympatric and closely related butterflies, showing separated niches at the microhabitat level. *Pieris rapae* significantly selected O microhabitats to oviposit, while *P. napi* was more frequently detected at OC microhabitats. The larvae of *P. rapae* grew thus under completely different thermal conditions with marked interspecific differences in thermal exposure (in the range of 5–10°C). Fully matching these results, *P. rapae* presented greater tolerance to thermal stress than *P. napi*. Friberg & Wiklund (2019) recently found that these two butterfly species in nature selected different plant species on which to lay their eggs but had similar host preferences under laboratory conditions. These results suggested that habitat choice for oviposition could precede host-plant selection. Our study provides further support to the relevant role that habitat structure can have in oviposition behaviour. The two host-plant species we studied were present in all microhabitats, but both butterfly species oviposited more frequently on the host plants in their preferred microhabitats (Fig. 5, Table 2). We hypothesize that ovipositing females possibly use multiple cues following a spatially structured and hierarchical process: firstly, selecting a particular microhabitat in response to vegetation cues (e.g. vegetation closure or openness) and, subsequently, contrasting in the selected microhabitat plant-specific traits in order to choose between alternative individual plants (look at Friberg, Olofsson, Berger, Karlsson, & Wiklund, 2008; Friberg & Wiklund, 2019, for similar results). In our study, concurrent changes in host-plant and microclimatic conditions were produced by microhabitat modification. Either background vegetation, temperature, or host-plant traits associated with quality (such as foliar chlorophyll and water contents) can influence oviposition behaviour in insects (Gibbs & Van Dyck, 2009; Myers, 1985; Wolfson, 1980). Further experimental studies are therefore required to quantify how these diverse factors may sequentially influence oviposition decisions.

Our study highlights the importance of analysing variation at these finer scales, as recently indicated (De Frenne et al., 2013, 2019; Pincebourde et al., 2016; Suggitt et al., 2018; Woods et al., 2015, among others). Multiple and interacting fine-scale processes can simultaneously operate and modulate the ecological responses of insects to global stressors (Carnicer et al., 2017). For example, variation in butterfly phenology, host-plant species, microhabitat, and oviposition behaviour in a study of populations of *Euphydryas editha* (Nymphalidea, Lepidoptera) in California and southern Oregon produced a geographic mosaic of varying microclimates and thermal exposures. The spatiotemporal exploitation of this microclimatic mosaic could therefore potentially confer higher resilience to climate change in this climatically sensitive species. The site-specific interactions between microclimatic and host-plant variation across the open–closed ecotones in our study could also have key roles in determining the vulnerabilities of the butterfly populations to global warming and the increasing impacts of drought. *Pieris napi* abundance has decreased in the last two decades by more than an order of magnitude at the lowland site 2 associated with summer rainfall, but has moderately increased at the mid-elevation site (Carnicer et al., 2019). The main host plant of *P. napi* senesced at the lowland site during the period of maximum abundance (i.e. the second generation, in June), which triggered a marked deterioration of host-plant traits associated with quality (Fig. 1J, L, N). The host plants were then scarce for 2–3 weeks until fresh leaves from new shoots emerged from resprouting rhizomes (Fig. S4). Resprout densities, with more leaves and chlorophyll, however, were higher in the lowland O microhabitats, where *P. napi* rarely oviposited (Figs. 3, 4EE, and 5). Summer resprouts in OC microhabitats, which were more frequently selected by *P. napi*, had higher foliar water contents and less thermal stress but low densities and fewer leaves (Figs. 3, 4G, W, and 5). These results contrast with the scenario at the mid-elevation site, where high-quality host plants were available throughout the year, especially in the C and OC microhabitats that benefited from an effective buffering (Figs. 1 and 2). The yearly reduction at the lowland site of *P. napi* records between consecutive generations after peak abundance may be associated with the low availability of host plants in the microhabitat selected for laying eggs. The presence of alternative host-plant species (e.g. *Brassica nigra*) in a fresh stage should be discarded before confirming this hypothesis. Whether the *P. napi* decadal declines at the lowland site are caused by an increasing spatial discordance between *P. napi*, host plant and thermal exposure across ecotones during summer dry periods remains to be assessed, however.

The increased impacts of drought linked to climate change and habitat loss in the northwestern Mediterranean Basin have been negatively associated with the richness (Carnicer et al., 2013; Stefanescu, Carnicer, & Peñuelas, 2011; Stefanescu, Herrando, & Páramo, 2004), distribution (Wilson et al., 2005; Wilson, Gutiérrez, Gutiérrez, & Monserrat, 2007), and demographic trends (Carnicer et al., 2019; Herrando et al., 2019; Melero, Stefanescu, & Pino, 2016; Stefanescu, Torre, Jubany, & Páramo, 2011; Ubach, Páramo, Gutiérrez, & Stefanescu, 2019) of butterfly species. The abandonment of traditional land uses in recent decades has profoundly modified Mediterranean landscapes, promoting substantial forest expansion at the expense of semi-natural grassland and scrub (Debussche, Lepart, & Dervieux, 1999). These land-cover trends represent an important threat to most of the butterfly species in Catalonia, because about 90% of species are more strongly associated with open habitats (Ubach et al., 2019). Indeed, previous studies have found that vegetation encroachment was the cause of larger population declines and the more frequent local extinctions of open-habitat butterflies (Herrando et al., 2019; Melero et al., 2016; Stefanescu, Torre, et al., 2011; Ubach et al., 2019). An increase in the dominance of closed-habitat species has consistently been reported in many butterfly communities, especially in the warmer areas of Catalonia (Ubach et al., 2019). We have demonstrated that open habitats can attain temperatures that could be limiting for more thermosensitive species. Vegetation encroachment in warm areas might thus benefit them to the detriment of thermotolerant species, which are likely adapted to these warmer and open conditions. Ubach et al. (2019) also reported that shifts in butterfly communities towards higher frequencies of closed-habitat species were less marked in heterogeneous landscapes. Our results indicate that the variety of microhabitats that arise in landscape mosaics is associated with both biotic and abiotic variation, permitting the co-occurrence of two closely related species with similar host-plant use but separated niches at the microhabitat scale. The preservation of landscape mosaics, avoiding excessive vegetation encroachment, can therefore be a determinant of insect conservation during global change.

Numerous fingerprints of the effects of global climate change have already been identified in insect populations (Boggs, 2016). Most of the predictive models of the responses of organisms to climate change are based on correlative approaches that use coarse-grain data. However, several studies have evidenced the need to adopt a more mechanistic approach incorporating the processes that generate microclimatic variation at fine-scales ultimately determining the climatic exposure of organisms (Pincebourde et al., 2016; Suggitt et al., 2018; Woods et al., 2015). The case documented in our study adds strong support to this idea. Furthermore, it shows that predictive models including variability at the microhabitat scale should not only consider microclimatic variability but also other biotic and abiotic factors that may vary concurrently, buffering or exacerbating large-scale stressors.

## ACKNOWLEDGEMENTS

Emili Bassols and Francesc Xavier Santaeufemia provided support with permission management, scientific advice and key assistance during field work. Meritxell Garcia, Armand Casadó, Carlos López, Sofía Cortizas, Joaquim de Gispert, Katarzyna Bartnik and Agnieszka Juszczak contributed to the field and experimental work. Consorci per a la Protecció i Gestió dels Espais Naturals del Delta del Llobregat, Parc Natural de la Zona Volcànica de la Garrotxa, and Parc Natural Aiguamolls de l’Empordà provided logistic support. This research was supported by the Spanish Ministry of Science and Innovation through a doctoral grant (FPU17/05869).

