## Supporting information for "Varying thermal exposure, host-plant traits and oviposition behaviour across vegetation ecotones"

**Text S1. Characterization of the host plant microhabitats from open – closed ecotones. Measurement and analysis of the vegetation cover and classification of microhabitats according to their degree of closure**.

We have termed *open – closed ecotones* the areas where open habitats contacts with closed habitats and the transition zones in between, which present diverse microhabitats differing in their degree of closure. The present study assessed whether variation at this microhabitat level influenced microclimatic conditions, the traits of the host plants growing across open – closed ecotones and the interactions they stablish with two butterfly species that oviposit on them. For this reason, we measured vegetation cover in each microsite of host plant monitoring, which represented the different microhabitats from open – closed ecotones. Measurements of vegetation cover considered, on the one hand, the canopy that shrubs and trees provided and, on the other, the ground cover by herbaceous plants, as both vegetal layers could influence microenvironmental conditions of the studied host plants (medium-size herbs). The measurements were repeated each monitoring day in order to additionally assess the temporal dynamics of vegetation cover in each microsite.

Shrubs and trees influence on each microsite was assessed through the measurement of the canopy closure (i.e. "the proportion of the sky hemisphere obscured by vegetation when viewed from a single point", Jennings, 1999). Canopy closure is supposed to be more tightly associated to light and microclimatic conditions at the understorey than canopy cover, which just takes into account the vertical projection of tree crowns on the floor (Jennings, 1999). The method of measurement consisted in visually estimating the per cent area occupied by the canopy assigning it to one of the cover classes defined by Daubenmire (1959) (0-5%, 5-25%, 25-50%, 50-75%, 75-95%, 95-100%) and taking its midpoint. The ocular estimation was conducted in each one of the vertical and the four cardinal directions, separating shrubs from trees. For the four cardinal directions, total canopy closure was then calculated as the mean of shrub closure and tree closure, as no overlapping between these two vegetal layers was assumed to occur. Because trees and shrub layers were considered to overlap in the vertical projection, the maximum of the two canopy closure values was taken as total canopy closure in this case. Finally, canopy closure in each microsite was calculated as the mean value of canopy closure in the five directions.

The herbaceous layer was characterized both as herbaceous ground cover and as mean herb height. Herbaceous cover was estimated by the point-intercept method. A rope with 15 knots separated by distances of 50 cm was randomly placed in the host plant microsite. For each knot, we annotated whether it contacted an herbaceous plant or, contrarily, if it had fallen on bare soil. Then, herbaceous cover was calculated as the relative frequency of knots covered by herbs. Additionally, herb plant height (cm) was also measured in the herbaceous plants contacted with the rope (or in the nearest herb in case the knot fell on bare soil) and the mean of the height of these 15 individuals was then calculated.

Data analyses were conducted with R version 3.6.1 (R Core Team, 2019). They were based on a careful examination of canopy closure, herbaceous cover and herbs height data including both analyses of spatial variation between host plant microsites in each study site (ANOVA and post-hoc Tukey test) and the assessment of their temporal dynamics (local polynomial regressions). Combining this information with field observations during the monitoring period, we finally classified the host plant microsites into four microhabitat types: closed (C), semi-closed (SC), semi-open (SO) and open (O). Photographs of different host plant microhabitat types for each site can be found in Figure S2. In Figure S3, the main results of vegetation cover analyses are shown. More open microhabitats (i.e. O and SO) were characterized by presenting a mean canopy closure of the whole monitoring period inferior than 50%. Because shrubs and trees had a sparser distribution there, the herbaceous layer was considered to be more determining for these microhabitats. Semi-open microhabitat categories were then assigned to open areas (such as fields or path margins) with taller herbs and higher herbaceous cover (Fig. S3). Microhabitats more integrated in the forested and shrubby areas of the open – closed ecotones presented values of mean canopy closure greater than 50%. Among them, those that tended to reach inferior values of maximum canopy closure (Fig. S3D) or reached them later (Fig. S3B) were classified as semi-closed microhabitats to differentiate them from the closed microhabitat category.

**Text S2. Implementation of the thermal death time experiments in *Pieris napi* and *Pieris rapae*.**

To assess whether *P. napi* and *P. rapae* show qualitatively different larval survival responses to thermal stress, we designed a static thermal death time experiment (Rezende, Castañeda, & Santos, 2014). Survival probability to a thermal stress is determined by both its intensity and duration (Rezende et al., 2014). Thermal death time experiments (TDT) allow predicting from first principles how environmental temperatures (considering the intensity and duration of the thermal exposure) can affect larval survival (Deutsch et al., 2008; Rezende et al., 2014). In this study, we implemented a thermal death time experiment in last instar larvae of *Pieris napi* and *P. rapae* with three static thermal treatments (40, 42 and 44 ºC).

The experiment was conducted on 223 individuals from 20 family lines collected at mid-elevation site 1 and lowland site 2 in summer of 2018 (Table S6). Females from both locations and species were captured and its offspring reared in growing chambers at 22 ºC 13L:11D, with fresh and abundant host plant (*Lepidium draba* and *Alliaria petiolata)*. Before the application of the thermal treatment, larvae were acclimated for 1 hour at constant 22ºC deprived of food. For the TDT experiment, we recorded the larval initial weights (g) and we subsequently placed the larvae in individual plastic pots (diameter: 3.5 cm, height: 7 cm) that were submerged in a water bath at a constant temperature (i.e. 40, 42 or 44ºC depending on the treatment). Their status (alive or dead) was checked at regular time interval periods (see Table S7 for details). Final larval weight (g) and time of death (min) were annotated. During the whole experimental treatment, the water temperature was continuously recorded using a thermal camera (FLIR-E90), and the air temperature inside the tube, with a temperature and humidity sensor (LascarElectronics EL-USB-2-LCD, 20-seconds resolution).

As observed in previous studies (Rezende et al., 2014), in each species, the time of death (min) and the temperature (ºC) of the TDT essays were significantly associated in semilogarithmic curves (Figure S5). TDT curves can be described as:

$$\log_{10} t = \frac{(\mathrm{CT}_{\max} - T)}{z}$$

where *t* is the observed time to death of the larvae in the $T_{ko}$ constant temperature applied, $CT_{max}$ is the temperature that would result in knockdown or death at 1 min ($\log_{10} t = 0$) and z is the constant of thermal susceptibility describing how thermal tolerance decays with the duration of the heat challenge (Rezende et al., 2014). From this equation, we estimated the upper limit of thermal tolerance ($CT_{max}$ ) and the thermal susceptibility (*z*, defined as the inverse of the slope of the TDT curve) of each species. Then, an analysis of the covariance (ANCOVA) modelling the effect of temperature, species and their interaction on larval death time was applied to assess whether the two species presented different slopes in the TDT curves. Additionally, general linear models were constructed for each species to evaluate the effect of thermal treatment (ºC), larval weight (g), site and family on $\log_{10}$ of larval death time. All data analyses were conducted with JMP (SAS Institute Inc., 2008).

Tables S8 and S9 summarize the results of the general linear models. Thermal treatment and larval weight were the most significant factors affecting larval death time of both species. The slopes of semilogarithmic relationships for *Pieris napi* and *P. rapae* were significantly different (ANCOVA test; p=0.0085, Figure S5). The estimates of thermal susceptibility *z* (*P. rapae* = 5.10 ± 0.24; *P. napi* = 4.10 ± 0.26) and $CT_{max}$ (*P. rapae* = 53.48ºC; *Pieris napi* = 51.08ºC) were higher in *Pieris rapae,* which indicated a greater tolerance to thermal stress (i.e. an inferior slope in the TDT curve).

As in Carnicer et al., 2019, the lethal thermal threshold with a daily exposure of 6 hours during the whole larval period (i.e. TE6H, an exposure of a total of 129h) was estimated from TDT equations. For *Pieris napi*, the critical temperature for a TE6h exposure was 35.14ºC and, for *P. rapae*, it was 33.65ºC. The mean between these two values, around 34.5ºC, was then taken as a reference temperature linked to stressful conditions affecting larval survival to compare it with thermal values recorded during the monitoring campaign of host plant growing microsites across open – closed ecotones.

**References from the supplementary materials**


Carnicer, J., Stefanescu, C., Vives-Ingla, M., López, C., Cortizas, S., Wheat, C. W., … Peñuelas, J. (2019). Phenotypic biomarkers of climatic impacts on declining insect populations: A key role for decadal drought, thermal buffering and amplification effects and host plant dynamics. *Journal of Animal Ecology*, *88*, 376–391. https://doi.org/10.1111/1365-2656.12933

Daubenmire, R. F. (1959). A canopy-coverage method of vegetational analysis. *Northwest Science*, *33*, 43–64.

Deutsch, C. A., Tewksbury, J. J., Huey, R. B., Sheldon, K. S., Ghalambor, C. K., Haak, D. C., & Martin, P. R. (2008). Impacts of climate warming on terrestrial ectotherms across latitude. *Proceedings of the National Academy of Sciences*, *105*(18), 6668–6672. https://doi.org/10.1073/pnas.0709472105

Jennings, S. (1999). Assessing forest canopies and understorey illumination: canopy closure, canopy cover and other measures. *Forestry*, *72*(1), 59–74. https://doi.org/10.1093/forestry/72.1.59

R Core Team. (2019). *R: A Language and Environment for Statistical Computing*. Retrieved from https://www.r-project.org/

Rezende, E. L., Castañeda, L. E., & Santos, M. (2014). Tolerance landscapes in thermal ecology. *Functional Ecology*, *28*(4), 799–809. https://doi.org/10.1111/1365-2435.12268

SAS Institute Inc. (2008). *JMP*. Retrieved from www.jmp.com

**Table S1. Census of oviposition.** **C**ensus effort (estimated as the total time devoted to census) and number of observations are summarised for each study site. O: open microhabitat, OC: intermediate microhabitat, and C: closed microhabitat.

| Location | Microhabitat | Census duration (min) | *P. napi* eggs (females) | *P. rapae* eggs (females) |
|---|---|---|---|---|
| **Mid-elevation population** | C | 579 | 3 (3) | 0 (0) |
| | OC | 924 | 46 (11) | 0 (0) |
| | O | 494 | 4 (3) | 10 (5) |
| | **TOTAL** | **1997** | **53 (17)** | **10 (5)** |
| **Lowland population** | C | 1895 | 0 (0) | 1 (1) |
| | OC | 2705 | 11 (10) | 3 (3) |
| | O | 620 | 0 (0) | 61 (7) |
| | **TOTAL** | **5220** | **11 (10)** | **65 (11)** |
| | | **7217** | **64 (27)** | **75 (16)** |

**Table S2. Temporal trends of microclimatic conditions and host plant traits.** Neighbourhood parameter α controlling the degree of smoothing of each local polynomial regression applied between the response variable and Julian day.

| Dimension | Response variable | Neighbourhood parameter (α) |
|---|---|---|
| **Microenvironmental conditions** | Leaf thermal amplification | 0,75 |
| | Soil humidity | 0,5 |
| **Host plant phenological, morphological and physiological traits** | Proportion of reproductive individuals | 0,65 |
| | Stem length | 0,75 |
| | Leaf water content | 1 |
| | Chlorophyll content | 1 |
| **Butterfly phenology** | Abundance index | 0,25 |

**Table S3. Flowering onset.** A summary of the fit of local polynomial regressions between the proportion of plants at reproductive stage and Julian day (neighbourhood parameter α = 0.5) in each microhabitat. RSE: residual standard error.

| Location | Microhabitat | Number of days | RSE |
|---|---|---|---|
| **Lowland population** | Closed (C) | 15 | 0,17 |
| | Semi-closed (SC) | 15 | 0,20 |
| | Semi-open (SO) | 15 | 0,19 |
| | Open (O) | 15 | 0,16 |
| **Mid-elevation population** | Closed (C) | 15 | 0,00 |
| | Semi-closed (SC) | 13 | 0,02 |
| | Semi-open (SO) | 16 | 0,14 |
| | Open (O) | 16 | 0,10 |

**Table S4. Microclimatic and host plant variation across open – closed ecotones**. A summary of the results from the ANOVA tests between the response variable and microhabitat type (C-SC-SO-O).

| Dimension | Response variable | Mid-elevation site (*Alliaria petiolata*) | | | Lowland site (*Lepidium draba*) | | |
|---|---|---|---|---|---|---|---|
| | | n | p-value | $R^2$ | n | p-value | $R^2$ |
| Microclimate | Leaf temperature | 567 | <0,0001 | 0,070 | 562 | <0,0001 | 0,383 |
| | Leaf thermal amplification | 567 | <0,0001 | 0,275 | 562 | <0,0001 | 0,425 |
| | Soil Humidity | 259 | <0,0001 | 0,190 | 383 | <0,0001 | 0,162 |
| Host plant morphological and physiological traits | Stem length | 40 | 0,0005 | 0,384 | 40 | <0,0001 | 0,713 |
| | Leaf length | 376 | <0,0001 | 0,132 | 489 | <0,0001 | 0,052 |
| | Leaf water content | 234 | <0,0001 | 0,179 | 251 | <0,0001 | 0,308 |
| | Leaf density | 326 | <0,0001 | 0,114 | 335 | <0,0001 | 0,318 |
| | Leaf chlorophyll content | 285 | 0,0004 | 0,062 | 403 | <0,0001 | 0,489 |

**Table S5. Seasonal variation of microclimatic and host plant traits patterns across open – closed ecotones.** A summary of the results from the ANOVA tests between the response variable and microhabitat type (C-SC-SO-O). Bold values indicate p-values below the selected level of significance (α = 0.05).

| | Response variable | Spring plants (Mar - May) | | | Senescent plants (Jun) | | | Summer new shoots (Jul - Aug) | | | Autumn new shoots (Sep - Oct) | | |
|---|---|---|---|---|---|---|---|---|---|---|---|---|---|
| | | n | p-value | $R^2$ | n | p-value | $R^2$ | n | p-value | $R^2$ | n | p-value | $R^2$ |
| Mid-elevation site (*Alliaria petiolata*) | Leaf temperature | 288 | **<0,0001** | 0,27 | 89 | **<0,0001** | 0,67 | 110 | **<0,0001** | 0,24 | 80 | **<0,0001** | 0,37 |
| | Leaf thermal amplification | 288 | **<0,0001** | 0,39 | 89 | **<0,0001** | 0,67 | 110 | **<0,0001** | 0,35 | 80 | **<0,0001** | 0,34 |
| | Soil humidity | 96 | **<0,0001** | 0,49 | 61 | **<0,0001** | 0,46 | 69 | **0,0002** | 0,28 | 33 | **<0,0001** | 0,75 |
| | Stem length | 20 | 0,0661 | 0,35 | 15 | 0,0694 | 0,46 | 20 | - | - | 20 | - | - |
| | Leaf length | 186 | **<0,0001** | 0,17 | 44 | **0,0002** | 0,38 | 93 | **0,0289** | 0,10 | 53 | **0,0031** | 0,24 |
| | Leaf water content | 158 | **<0,0001** | 0,48 | 49 | 0,8876 | 0,01 | 79 | 0,3707 | 0,04 | 44 | 0,7097 | 0,03 |
| | Leaf density | 165 | **<0,0001** | 0,15 | 41 | 0,1817 | 0,12 | 76 | **0,0006** | 0,21 | 44 | **0,0160** | 0,23 |
| | Leaf chlorophyll content | 186 | **0,0003** | 0,07 | 58 | **0,0003** | 0,31 | 99 | **<0,0001** | 0,19 | 53 | 0,1153 | 0,11 |
| Lowland site (*Lepidium draba*) | Leaf temperature | 252 | **<0,0001** | 0,43 | 68 | **<0,0001** | 0,75 | 176 | **<0,0001** | 0,52 | 66 | **<0,0001** | 0,49 |
| | Leaf thermal amplification | 252 | **<0,0001** | 0,48 | 68 | **<0,0001** | 0,75 | 176 | **<0,0001** | 0,40 | 66 | **<0,0001** | 0,47 |
| | Soil humidity | 84 | **<0,0001** | 0,35 | 65 | **<0,0001** | 0,58 | 168 | **<0,0001** | 0,37 | 66 | 0,4469 | 0,04 |
| | Stem length | 20 | **<0,0001** | 0,88 | 9 | **0,0119** | 0,77 | 20 | - | - | 20 | - | - |
| | Leaf length | 219 | **<0,0001** | 0,10 | 42 | **0,0019** | 0,32 | 162 | **<0,0001** | 0,15 | 66 | **0,0019** | 0,21 |
| | Leaf water content | 177 | **<0,0001** | 0,24 | 48 | 0,1191 | 0,16 | 54 | **<0,0001** | 0,51 | 57 | **<0,0001** | 0,35 |
| | Leaf density | 188 | **<0,0001** | 0,47 | 36 | **0,0001** | 0,48 | 54 | **0,0005** | 0,26 | 57 | **0,0009** | 0,27 |
| | Leaf chlorophyll content | 219 | **<0,0001** | 0,37 | 54 | **0,0159** | 0,25 | 164 | **0,0123** | 0,22 | 66 | **0,0011** | 0,28 |

**Table S6.** Number of larvae used in the thermal death experiments (TDT) for each static thermal treatment (44, 42 and 40 ºC), site, species and family line.

| | **Total** | **44 ºC** | **42 ºC** | **40 ºC** |
|---|---|---|---|---|
| **Mid-elevation site 1** | **103** | **45** | **38** | **20** |
| *P. napi* (4 families) | *31* | *15* | *16* | *0* |
| Female 1 (PN2) | 7 | 7 | 0 | 0 |
| Female 2 (PN22) | 6 | 2 | 4 | 0 |
| Female 3 (PN23) | 3 | 0 | 3 | 0 |
| Female 4 (PN24) | 15 | 6 | 9 | 0 |
| *P. rapae* (5 families) | *72* | *30* | *22* | *20* |
| Female 1 (PR10) | 8 | 8 | 0 | 0 |
| Female 2 (PR2) | 1 | 0 | 0 | 1 |
| Female 3 (PR22) | 19 | 6 | 7 | 6 |
| Female 4 (PR23) | 14 | 5 | 5 | 4 |
| Female 5 (PR25) | 30 | 11 | 10 | 9 |
| **Lowland site 2** | **120** | **40** | **40** | **40** |
| *P. napi* (6 families) | *60* | *20* | *20* | *20* |
| Female 1 (PNA) | 5 | 5 | 0 | 0 |
| Female 2 (PNB) | 27 | 5 | 5 | 17 |
| Female 3 (PNC) | 5 | 5 | 0 | 0 |
| Female 4 (PND | 10 | 5 | 5 | 0 |
| Female 5 (PNE) | 6 | 0 | 5 | 1 |
| Female 6 (PNF) | 7 | 0 | 5 | 2 |
| *P. rapae* (5 families) | *60* | *20* | *20* | *20* |
| Female 1 (PRA) | 15 | 5 | 5 | 5 |
| Female 2 (PRD) | 15 | 5 | 5 | 5 |
| Female 3 (PRE) | 5 | 0 | 5 | 0 |
| Female 4 (PRF) | 15 | 5 | 5 | 5 |
| Female 5 (PRG) | 10 | 5 | 0 | 5 |
| **Total (site 1 + site 2)** | **223** | **85** | **78** | **60** |

**Table S7. Thermal death time experiments**. Frequency of observation of the larval status (dead/alive) during the implementation of each thermal treatment.

| Thermal treatment | Measurement frequency |
|---|---|
| 40 ºC | Once every 30 min |
| 42 ºC | Once every 20 min |
| 44 ºC | Once every 10 min |

**Table S8. General linear model of larval death time of *Pieris napi***. Effect tests of thermal treatment (ºC), larval weight (g), site and family on $\log_{10}$ of death time (min) of *Pieris napi* larvae during TDT experiments. Model fit: $R^2$=0.81, p<0.0001, n=89 individuals.

| Source | DF | Sum of squares | F ratio | Prob > F |
|---|---|---|---|---|
| Thermal treatment (ºC) | 1 | 7.047 | 182.66 | <0.0001* |
| Larval weight (g) | 1 | 0.235 | 6.09 | 0.0158* |
| Site | 1 | 0.133 | 3.46 | 0.0668 |
| Family[site] | 8 | 0.867 | 2.81 | 0.0087* |

**Table S9. General linear model of larval death time of *Pieris rapae***. Effect tests of thermal treatment (ºC), larval weight (g), site and family on $\log_{10}$ of death time (min) of *Pieris rapae* larvae during TDT experiments. Model fit: $R^2$=0.82, p<0.0001, n=121 individuals.

| Source | DF | Sum of squares | F ratio | Prob > F |
|---|---|---|---|---|
| Thermal treatment (ºC) | 1 | 12.66 | 424.01 | <0.0001* |
| Larval weight (g) | 1 | 0.47 | 15.65 | 0.0001* |
| Site | 1 | $8.81e^{-6}$ | $3e^{-4}$ | 0.9863 |
| Family[site] | 8 | 0.30 | 1.2543 | 0.2750 |

**Figure S1. Geographic distribution of the studied areas.** A: Protected Natural Parks from Catalonia (NE Spain, inner panel). The mid-elevation site 1 is located in La Garrotxa Volcanic Zone Natural Park (green) and the lowland site 2 is located in Aiguamolls de l'Empordà Natural Park (orange). Grey areas correspond to the other existing Natural Parks. B, C: Location of the two monitored cohorts of host plants (dark red point) and the nearest meteorological stations (black point) in site 1 (panel B) and site 2 (panel C).

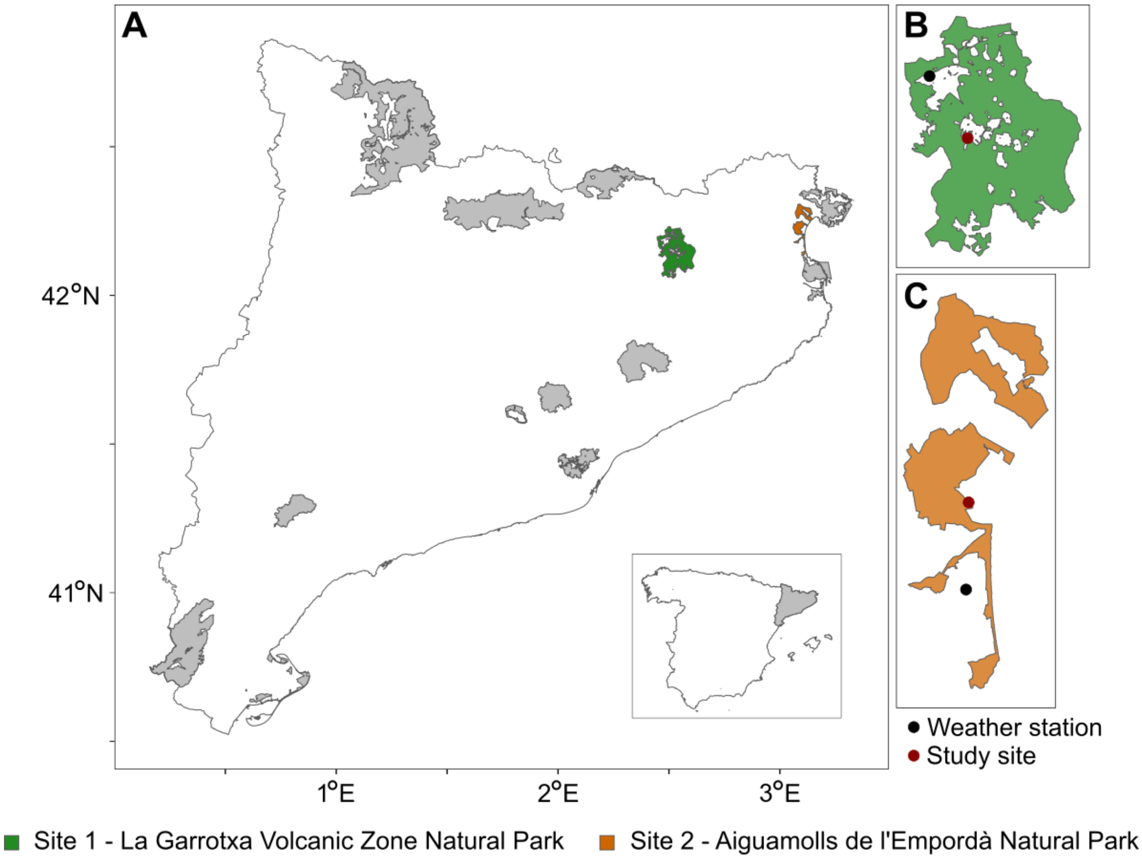

**Figure S2. Photographs of the landscape mosaic of the two study sites and the different microhabitat types of host plant growth present in open – closed ecotones.**

Mid-elevation site 1. La Garrotxa Volcanic Zone Natural Park. Landscape view (October).

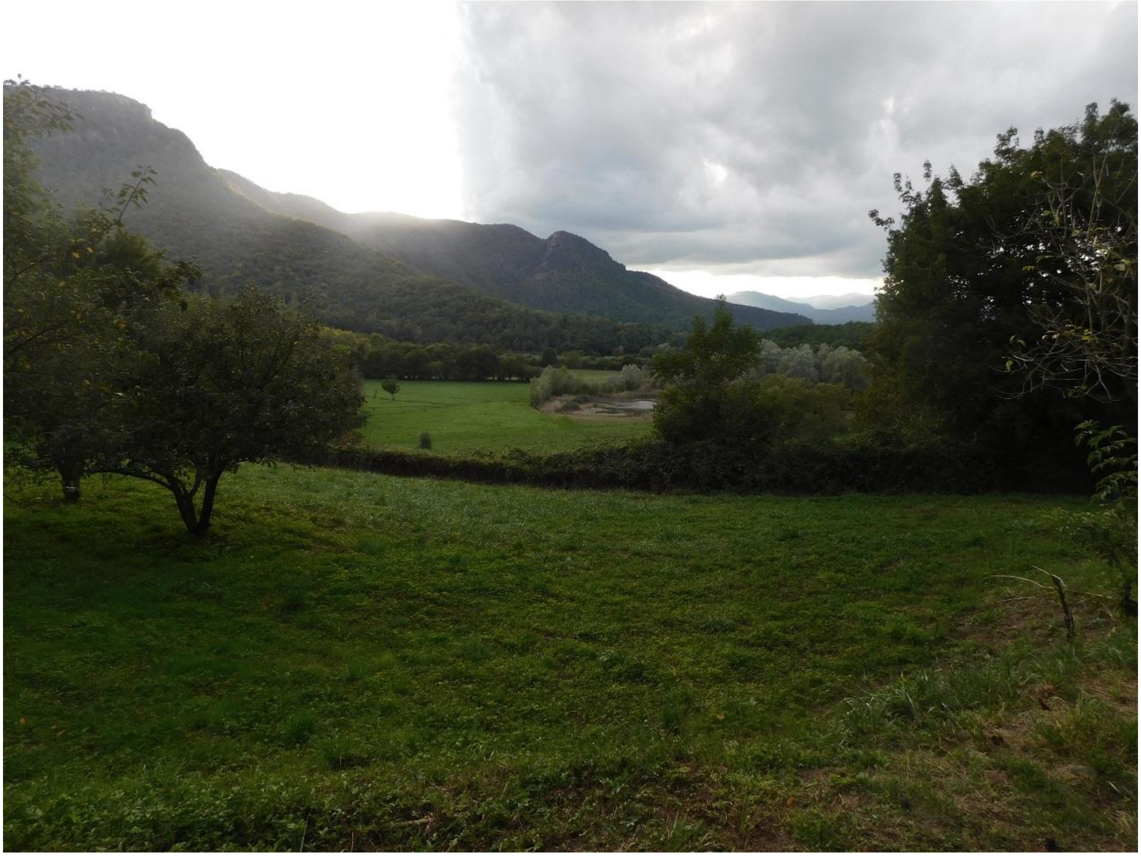

Closed microhabitat from mid-elevation site 1 during October.

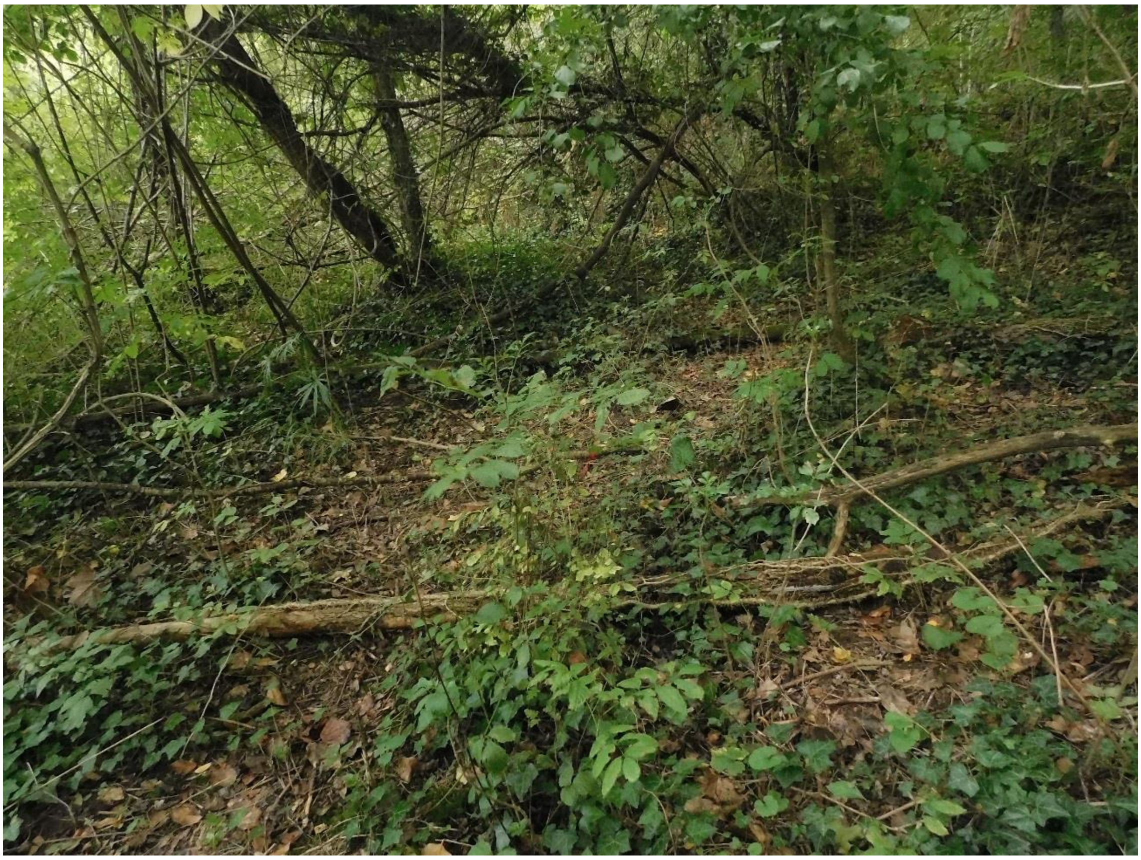

Semi-closed microhabitat from the mid-elevation site 1 during October.

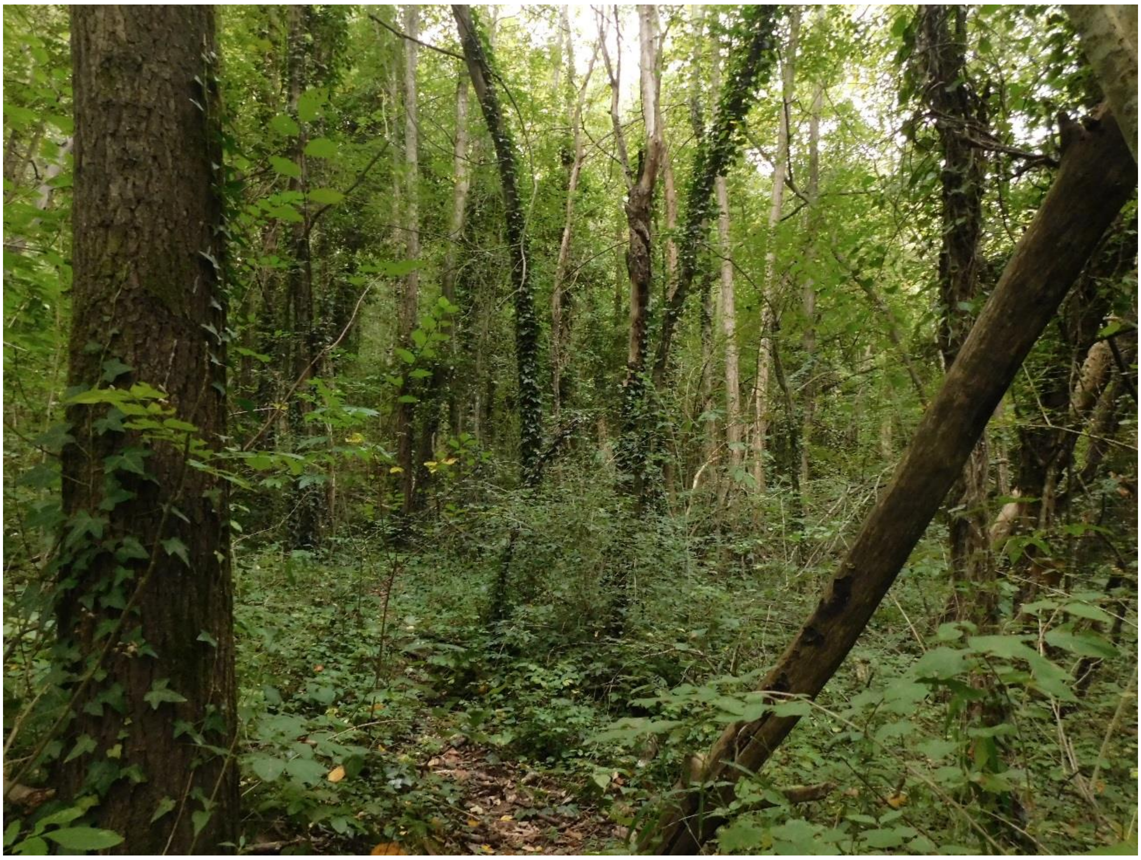

Semi-open microhabitat from the mid-elevation site 1 during October. The monitoring campaign was conducted before the fence that can be observed in the photograph was placed, when tall herbs like those that appear in the right side of the fence covered a much greater area.

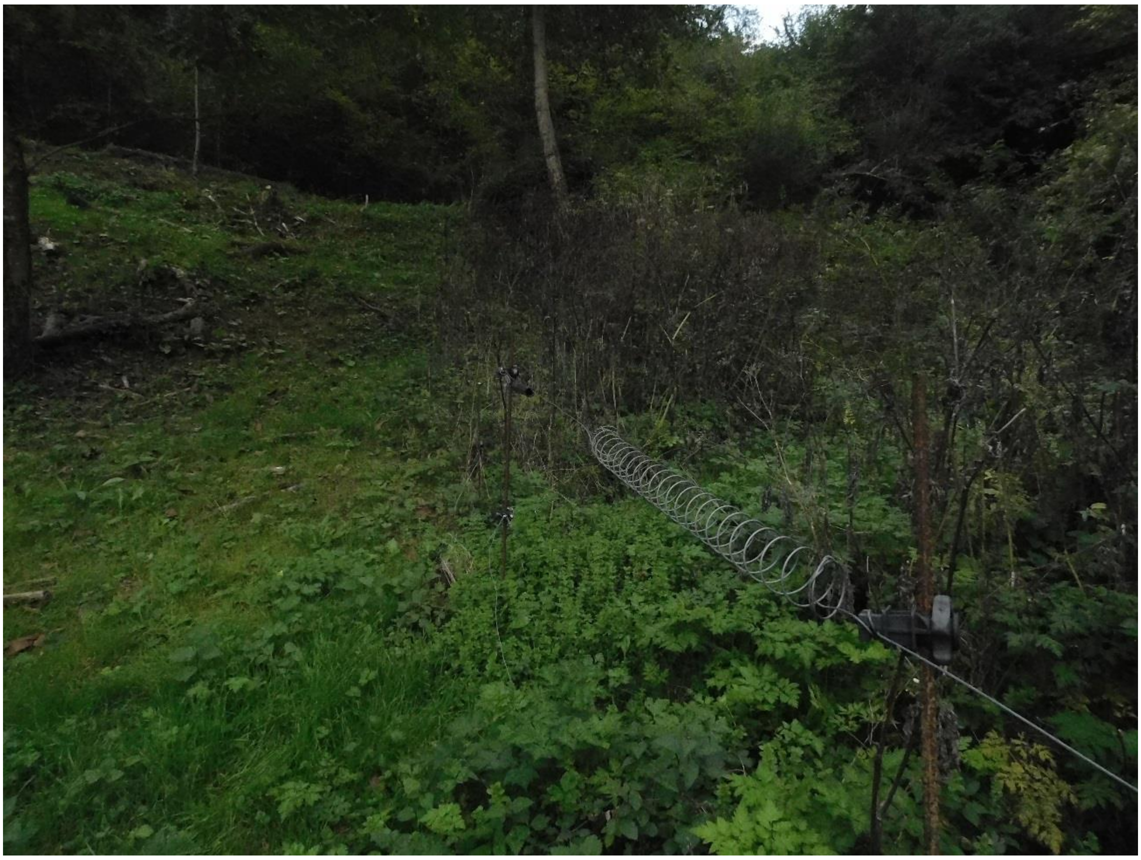

Open microhabitat from the mid-elevation site 1 during October. In the ecotone between the forest margin and the path, shrubs and trees covered less than 50% of the microhabitat and ground was sparsely covered by short and medium-sized herbs exposing *Alliaria petiolata* to sunlight conditions.

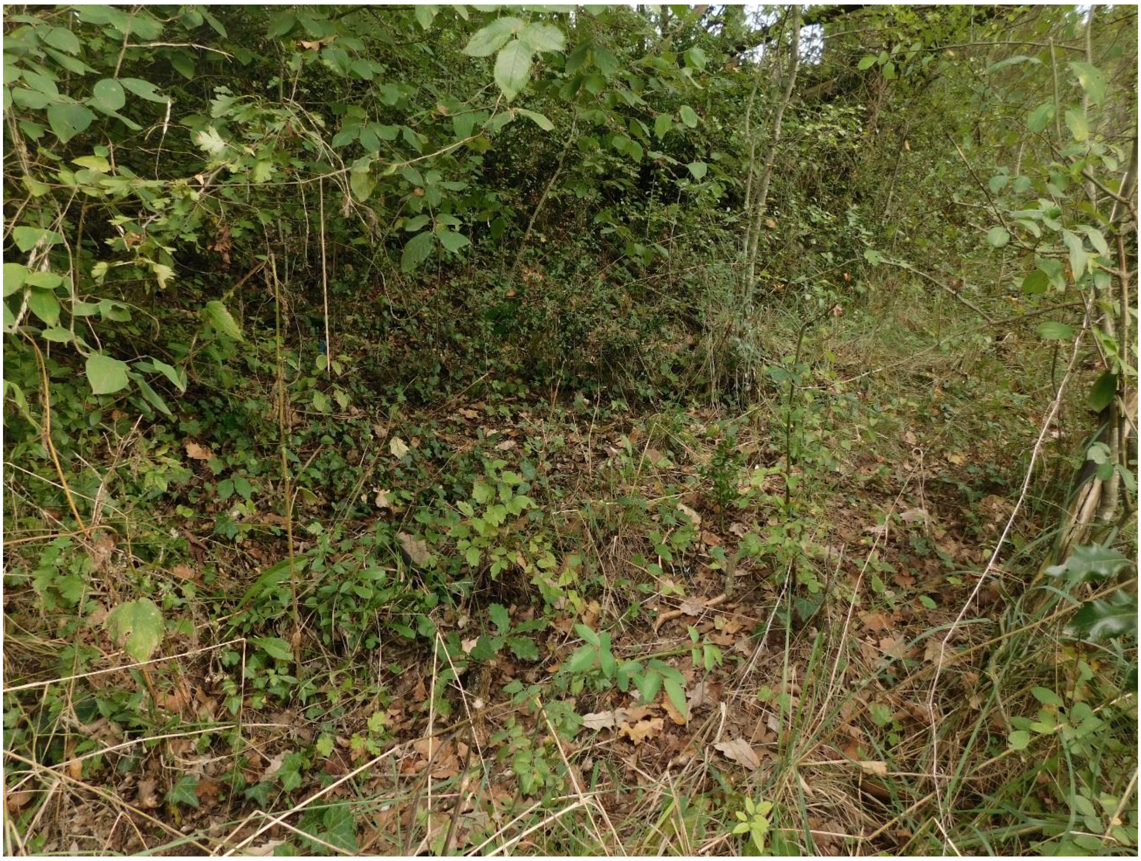

Non-flowering first-year rosettes of *Alliaria petiolata* that persisted in the mid-elevation site 1 after the fructification and senescence of the second-year individuals. The photograph has been taken during October. In the next spring, during the growing season, these rosettes will continue and complete their life cycle.

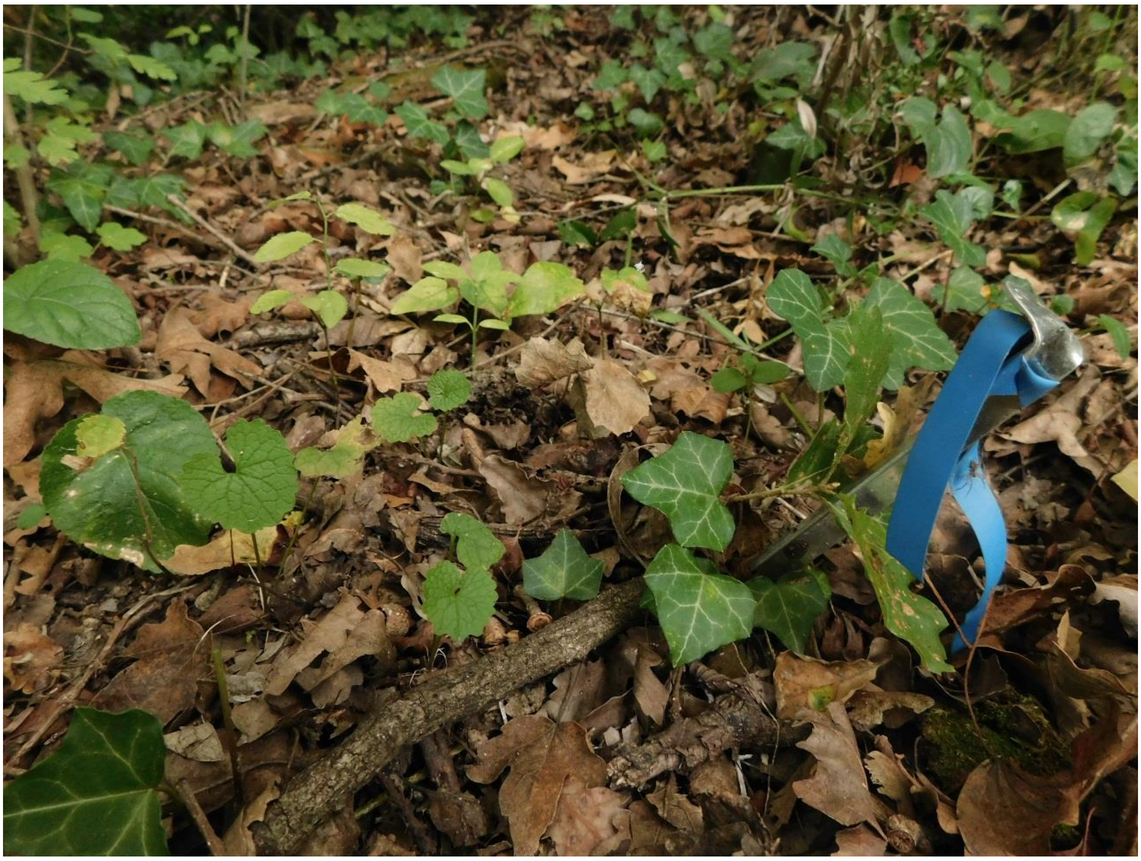

Lowland site 2. Aiguamolls de l’Empordà Natural Park. Landscape view. Early spring (March)

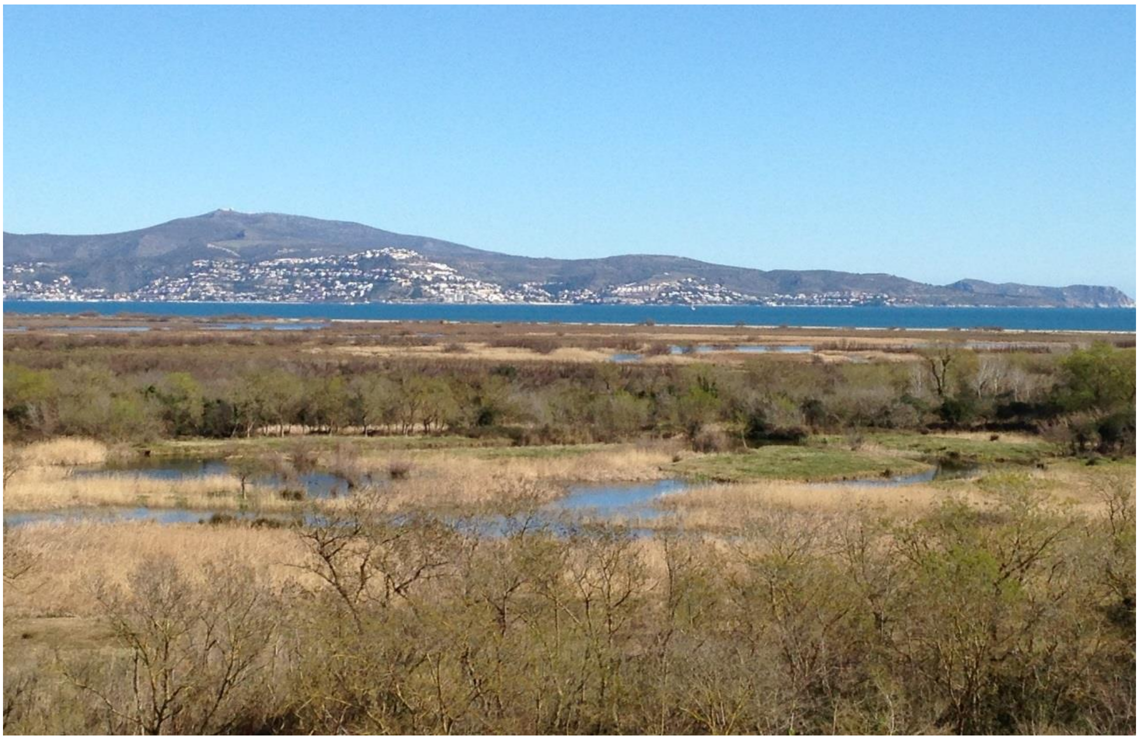

Closed microhabitat from the lowland site 2 during June. Senescent individuals of the host plant *Lepidium draba* could be detected in the ground before achieving a reproductive stage.

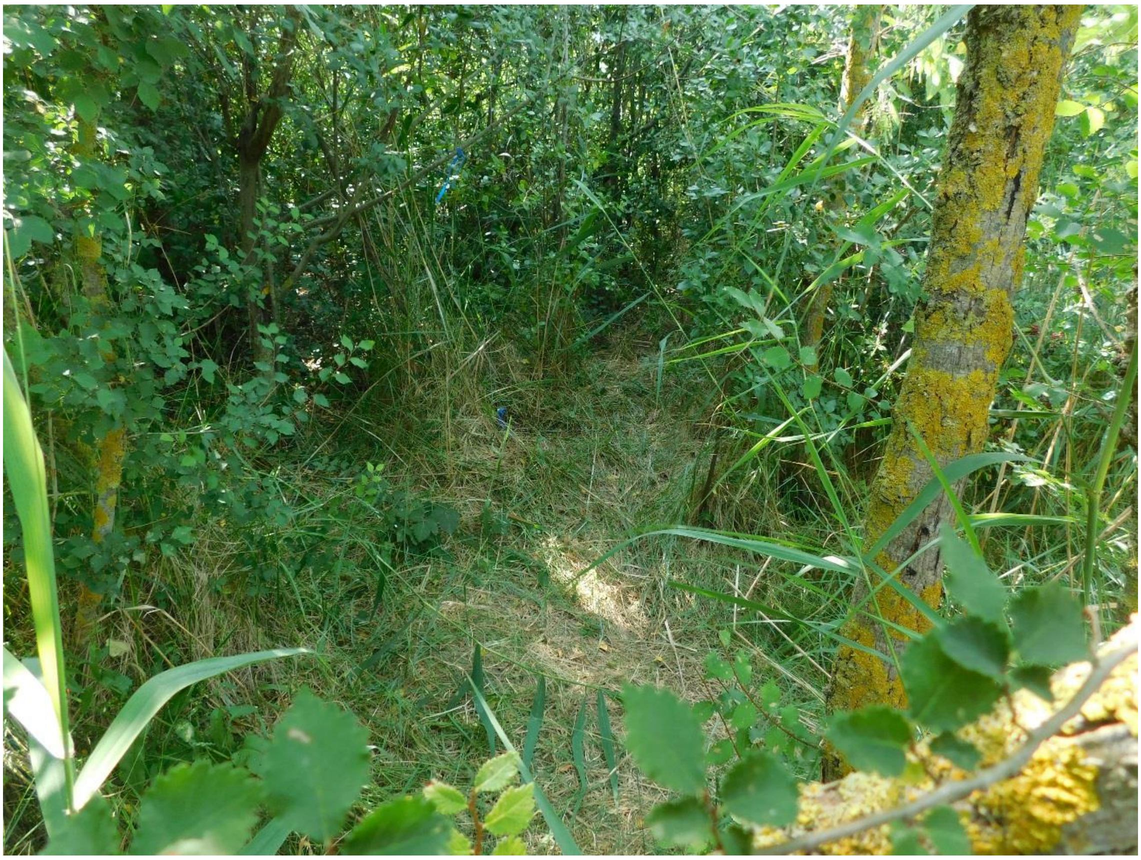

Semi-closed microhabitat from the lowland site 2 during May. Low light microconditions inhibited sexual maturation of *Lepidium draba* individuals, which presented an unusual vine-like growth form.

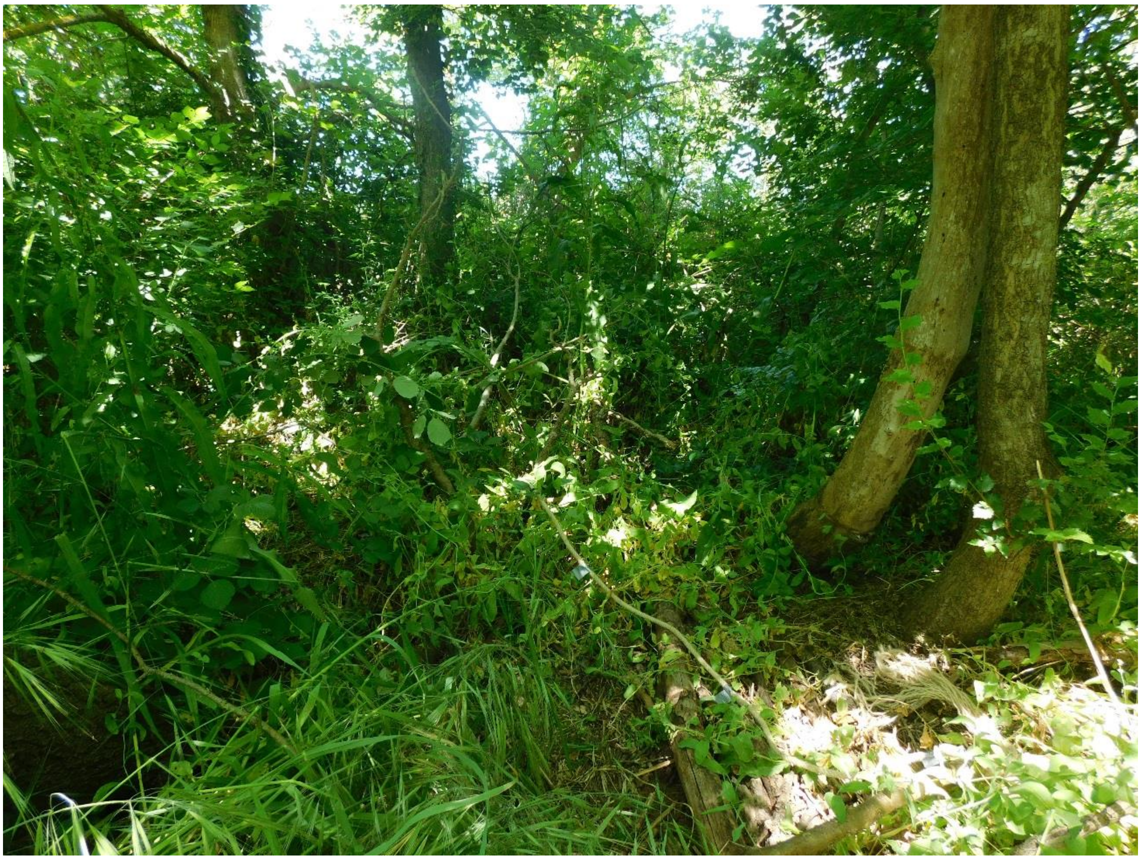

Semi-open microhabitat of the lowland site 2 during March. *Lepidium draba* still presented a vegetative stage.

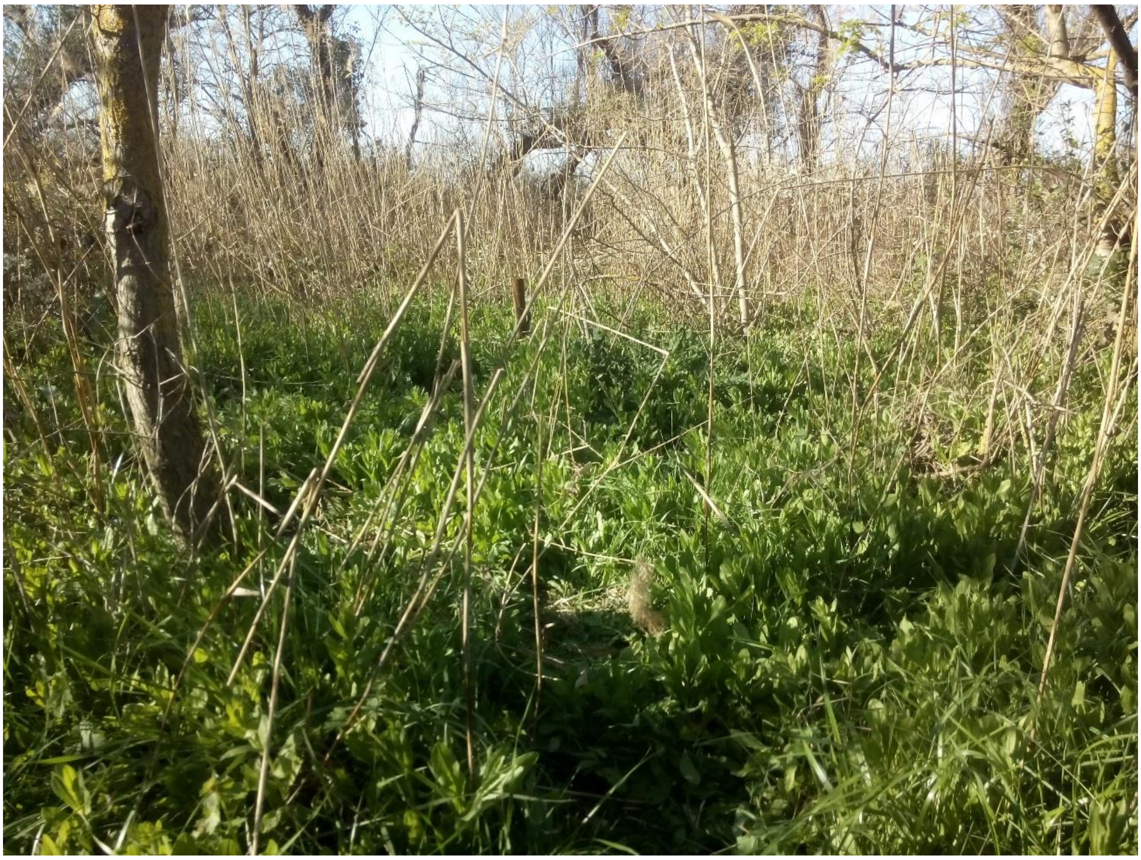

Open microhabitat from the lowland site 2 during May with *Lepidium draba* individuals at their fruiting stage.

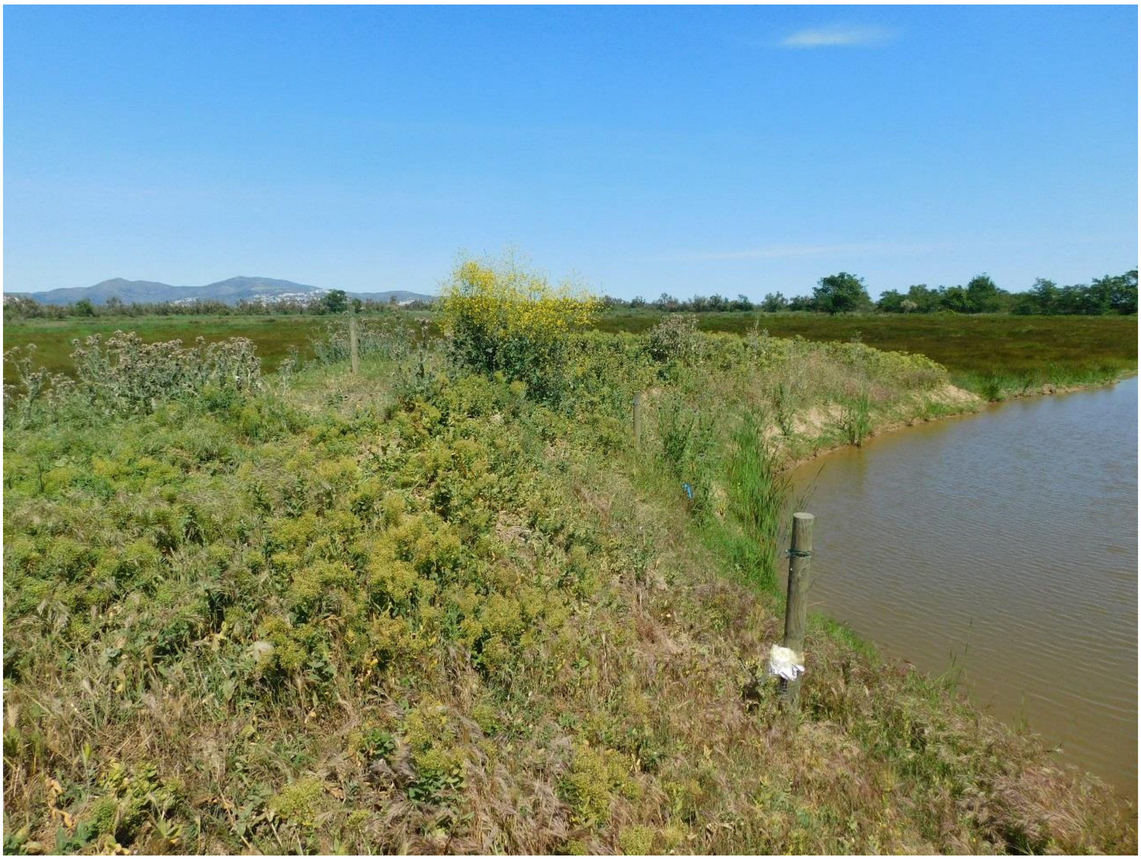

*Lepidium draba* resprouts emerging from subterranean rhizomes during midsummer (August) in an open microhabitat from the lowland site 2.

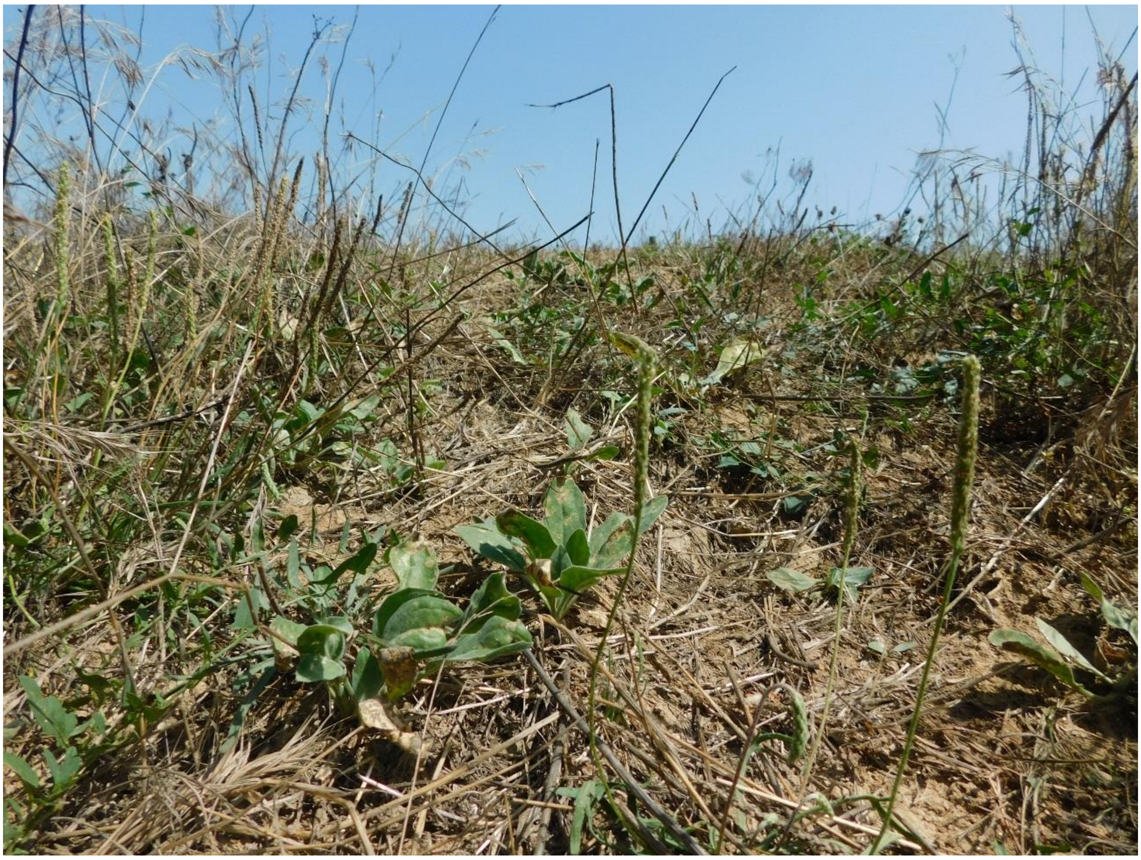

**Figure S3. Spatial and temporal variation of the vegetation cover in different microhabitat types defined in open – closed ecotones.** A, C: an ANOVA test comparing canopy closure percentage between closed and semi-closed microhabitats juxtaposed to an ANOVA test comparing mean herb height between semi-open and open microhabitats in the study sites of *Alliaria petiolata* (mid-elevation site 1, A) and *Lepidium draba* (lowland site 2, C). B, D: temporal dynamics of canopy closure (solid curves, C and SC microhabitats) and mean herb height (dashed curves, O and SO microhabitats) in the mid-elevation site 1 (B) and lowland site 2 (D). More details can be consulted in Text S1.

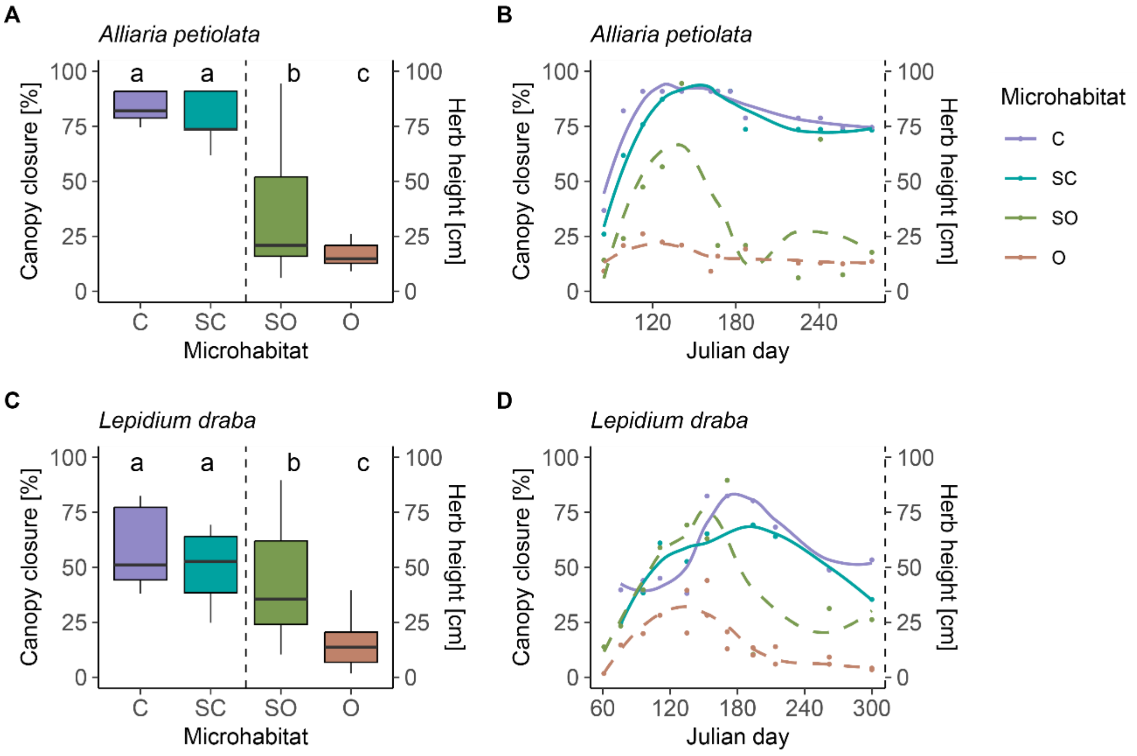

**Figure S4. Butterfly and host plant phenology in the two study sites.** Phenological progress of the host plant individuals growing in different microhabitats across the open – closed ecotones in the mid-elevation site 1 (A-D) and the lowland site 2 (F-I). Panels E and J: *Pieris napi* (black) and *Pieris rapae* (grey) phenologies obtained from the data of the transects of the Catalan Butterfly Monitoring Scheme located at both study sites. Ros: spring rosettes and young shoots before budding; rep: reproductive plants with buds, flowers and/or fruits; sen: senescent plants; summer ros: summer rosettes of *Alliaria petiolata* and midsummer resprouts of *Lepidium draba*.

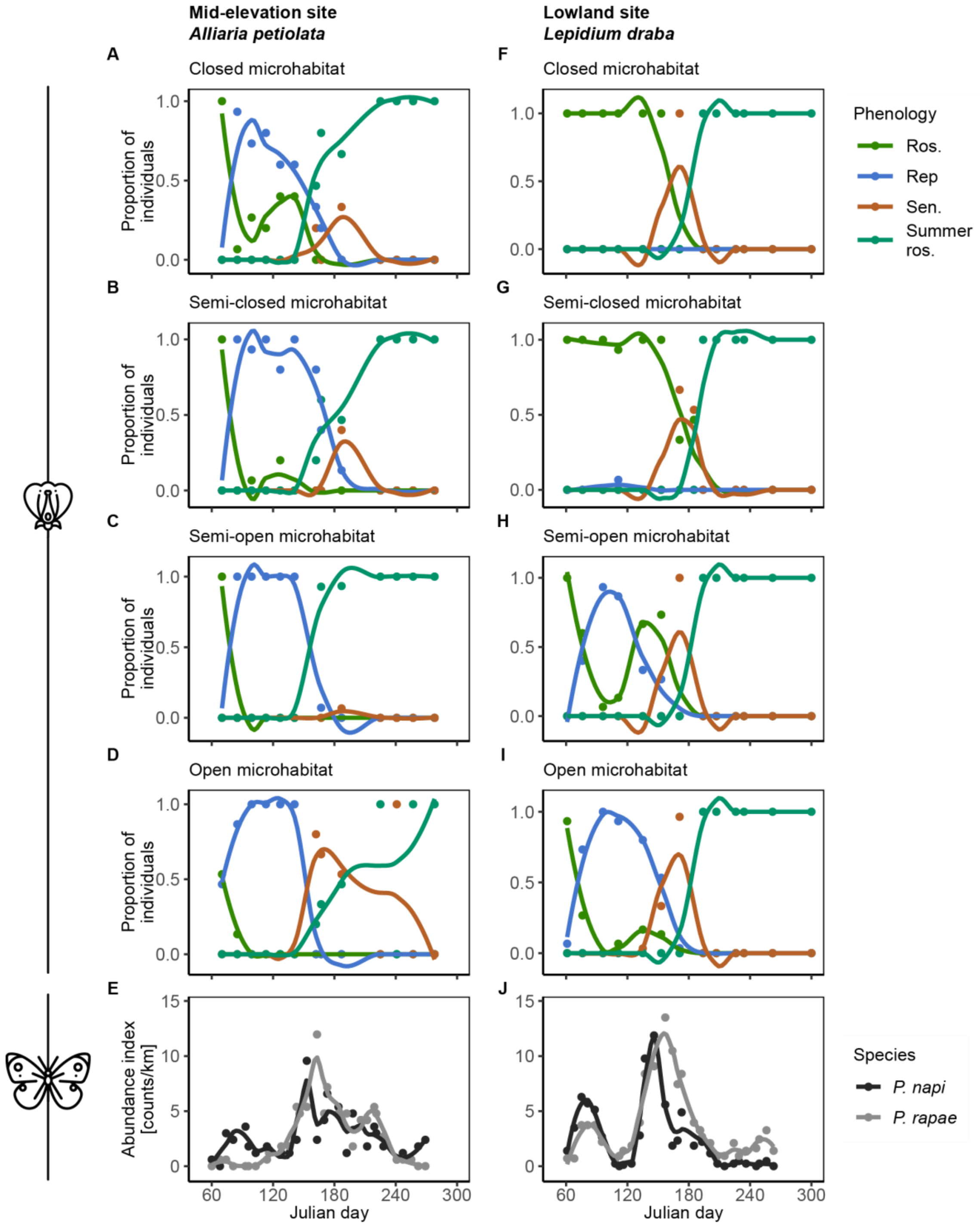

**Figure S5. A comparison of TDT curves for *Pieris napi* (blue) and *P. rapae* (red)**. Estimates of the thermal susceptibility constant (*z*), a test comparing the slopes (ANCOVA) and ordinary least squares fits for each species are reported.

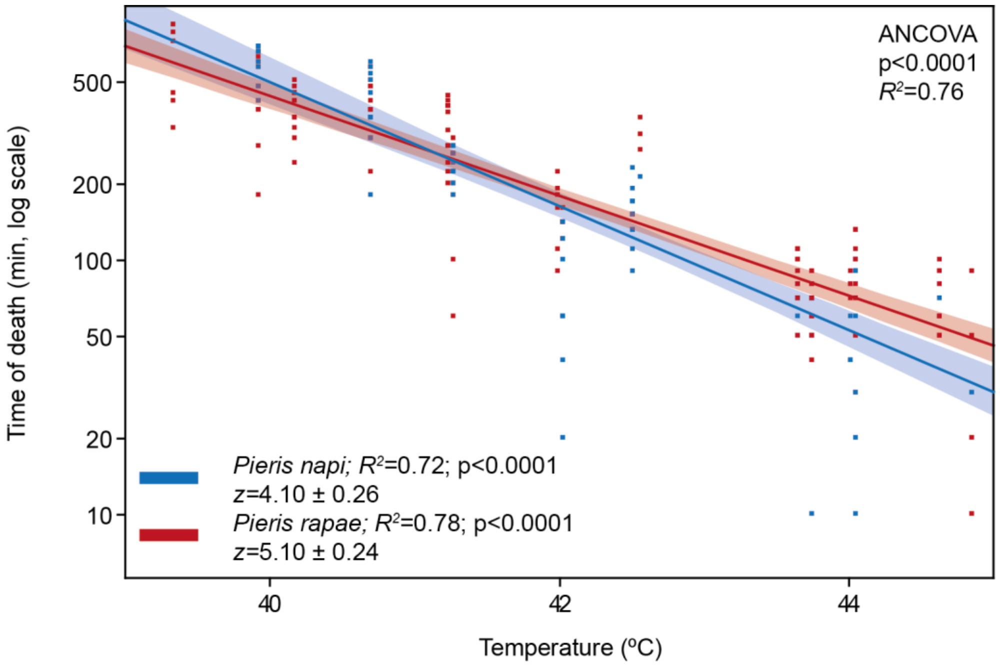

**Figure S6. Seasonal variation of the spatial patterns of variation across open – closed ecotones of the mid-elevation site 1.** Panels A-L: microclimatic conditions. Panels M-FF: *Alliaria petiolata* host plant traits. C: closed, SC: semi-closed, SO: semi-open, and O: open microhabitats. Different lower case letters indicate significant differences of the response variable between the corresponding microhabitat types in the Tukey HSD test. The dotted line in the panels A-D corresponds to the thermal threshold that would lead to larval death with a daily exposure of 6 hours during the whole developmental period (TE6H) estimated from the TDT experiments. The horizontal solid lines in the panels E-H indicate leaf temperatures equal to maximum daily air temperature. Positive values correspond to leaf thermal amplification phenomena, whereas negative values imply thermal buffer effects. A summary of the results is offered in Table S5.

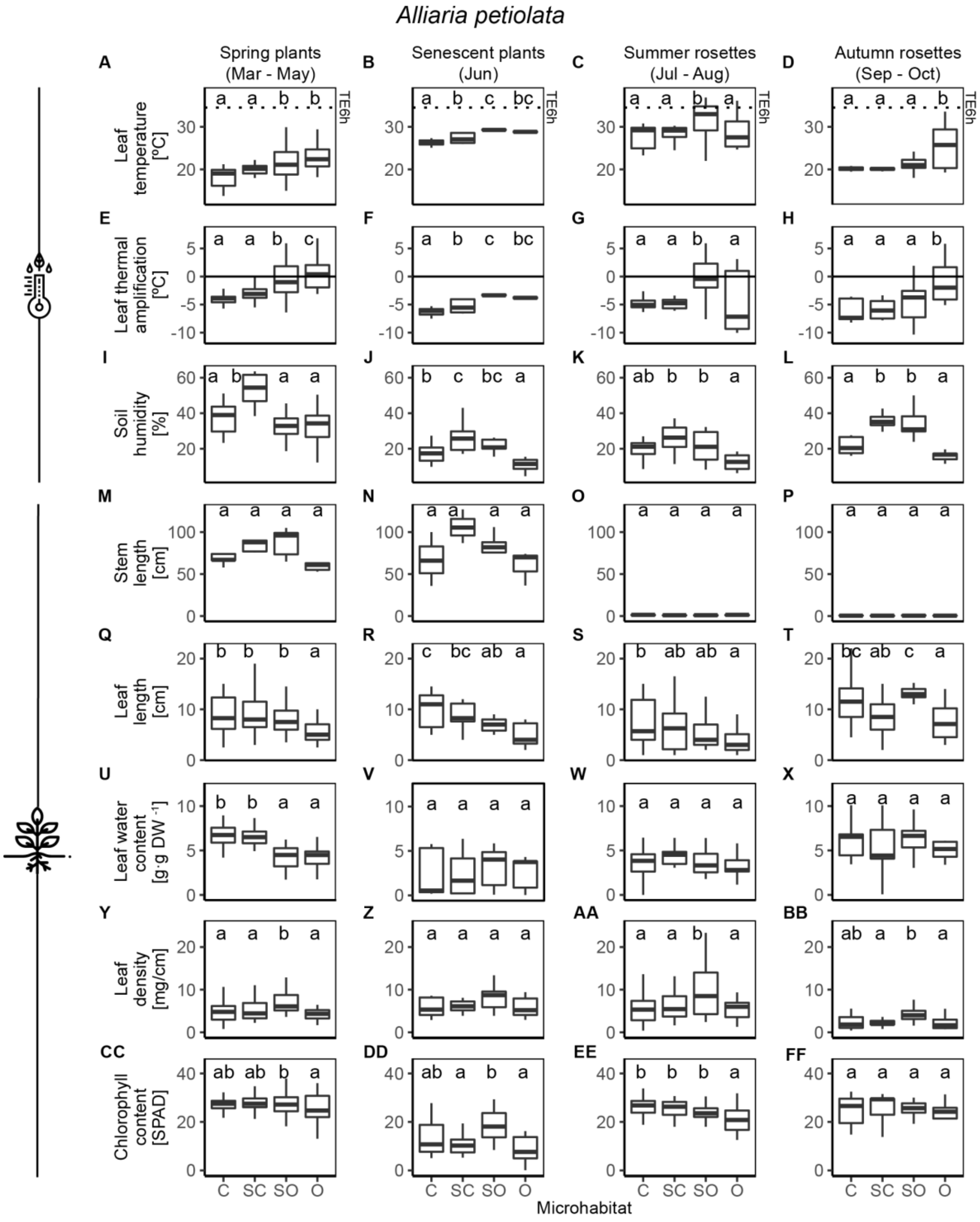
Alliaria petiolata
Spring plants (Mar - May)
Senescent plants (Jun)
Summer rosettes (Jul - Aug)
Autumn rosettes (Sep - Oct)
Leaf temperature [ºC]
Leaf thermal amplification [ºC]
Soil humidity [%]
Stem length [cm]
Leaf length [cm]
Leaf water content [g·g DW -1]
Leaf density [mg/cm]
Chlorophyll content [SPAD]
TE6h
C SC SO O
Microhabitat

**Figure S7. Seasonal variation of the spatial patterns of variation across open – closed ecotones of the lowland site 2.** Panels A-L: microclimatic conditions. Panels M-FF: *Lepidium draba* host plant traits. C: closed, SC: semi-closed, SO: semi-open, and O: open microhabitats. Different lower case letters indicate significant differences of the response variable between the corresponding microhabitat types in the Tukey HSD test. The dotted line in the panels A-D corresponds to the thermal threshold that would lead to larval death with a daily exposure of 6 hours during the whole developmental period (TE6H) estimated from the TDT experiments. The horizontal solid lines in the panels E-H indicate leaf temperatures equal to maximum daily temperature. Positive values correspond to leaf thermal amplification phenomena, whereas negative values imply thermal buffer effects. Because of the small number and size of midsummer resprouts in the semi-closed microhabitat, no leaf sample was taken to weight (panels W and AA) and leaf chlorophyll measurement was achieved in just one case (panel EE). A summary of the results is offered in Table S5.

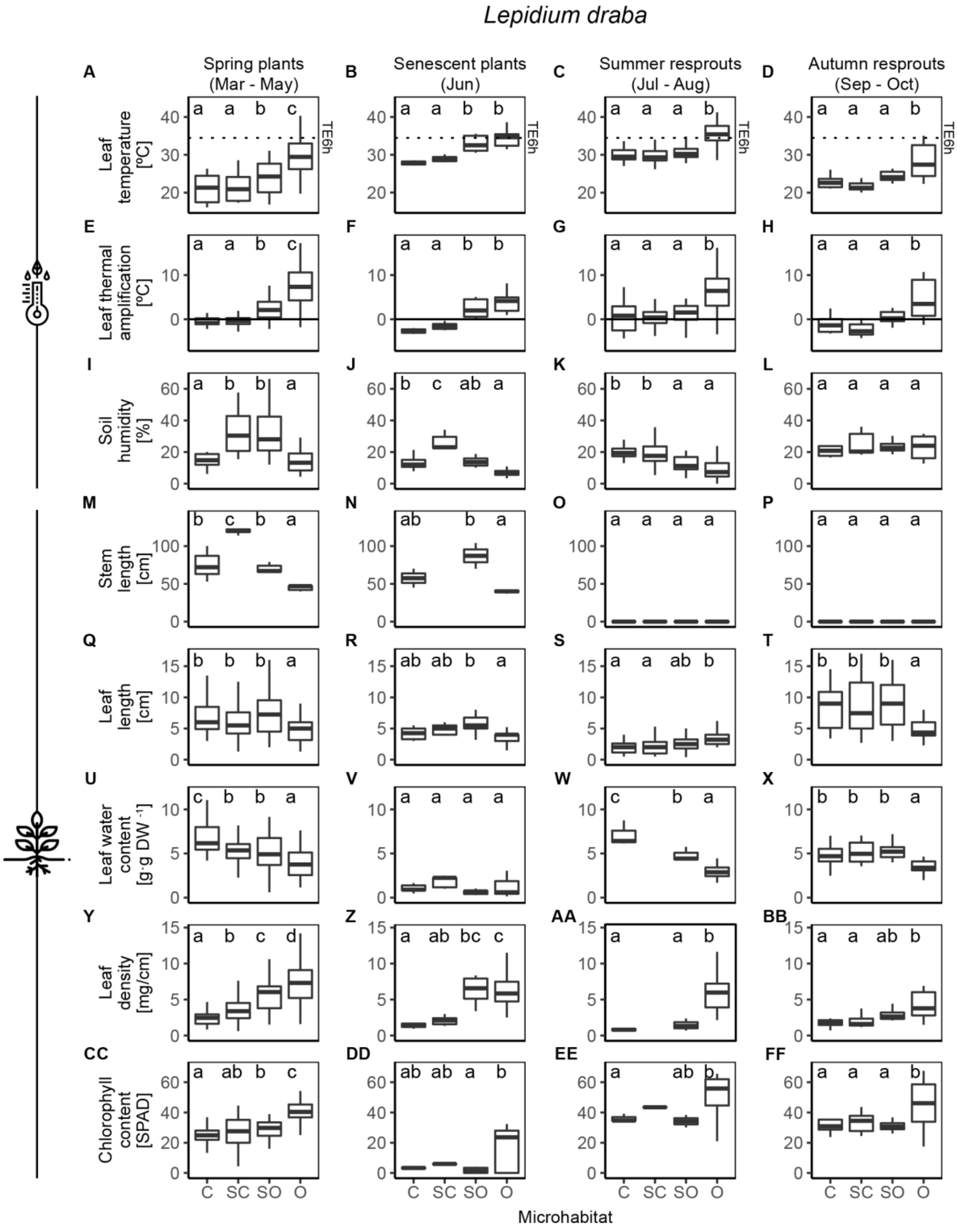
Lepidium draba
Spring plants (Mar - May)
Senescent plants (Jun)
Summer resprouts (Jul - Aug)
Autumn resprouts (Sep - Oct)
Leaf temperature [ºC]
Leaf thermal amplification [ºC]
Soil humidity [%]
Stem length [cm]
Leaf length [cm]
Leaf water content [g·g DW⁻¹]
Leaf density [mg/cm]
Chlorophyll content [SPAD]
TE6h
C SC SO O
Microhabitat
